# Aberrant retinal structure and vasculature in mouse models of dominant retinopathies caused by *CRX* homeodomain mutations

**DOI:** 10.64898/2026.03.19.712925

**Authors:** Chi Sun, Charles W. Pfeifer, Yiqiao Zheng, Rajendra S. Apte, Shiming Chen

## Abstract

CRX is a transcription factor essential for photoreceptor differentiation and functional development. Two missense mutations in CRX homeodomain, *CRX^E80A^* and *CRX^K88N^*, are linked to early-onset dominant retinopathies. Molecular studies have revealed distinct profiles of perturbed gene expression in differentiating photoreceptors of knock-in mouse models, resulting from altered DNA binding activities of mutant CRX proteins. This study characterizes concurrent retinal and vascular alterations in knock-in mouse models. Fated cones are present in heterozygous and homozygous *Crx^E80A^* and *Crx^K88N^* mutants at birth, but subsequent cone differentiation is rapidly compromised. Expression of rod marker rhodopsin (RHO) is absent in *Crx^K88N/N^*retinae but present in other mutants through adulthood. Notably, as compared to wildtype controls, RHO expression is prematurely activated in neonatal *Crx^E80A^* mutants. Among tested mutants, only *Crx^E80A/+^*retinae elaborate rod outer segments but still lose visual function by young adulthood. The presence of irregular retinal rosettes is a striking pathological phenotype in all mutants. Retinal rosettes displace the localization of inner neurons without affecting their cell numbers during retinal development. Retinal vessels develop close contact with rosette structures. In summary, disrupted photoreceptor differentiation leads to the loss of visual function and formation of retinal rosettes. The presence of retinal rosettes secondarily impairs the localization of inner neurons and vasculature. A deeper understanding of these cellular underpinnings will inform pathogenesis of CRX homeodomain mutations.

## INTRODUCTION

Inherited retinal diseases (IRDs) represent a group of genetic disorders characterized by progressive degeneration of photoreceptors. IRDs are caused by pathogenic gene mutations that are inherited in autosomal dominant, autosomal recessive, or X-linked patterns. Mutations in genes coding for photoreceptor-specific transcription factors, such as the *cone-rod homeobox* (*CRX*) gene, are associated with early-onset IRDs, notably Leber congenital amaurosis (LCA) and cone-rod dystrophy (CoRD)^1,2^. CRX is an essential transcription factor that regulates photoreceptor gene expression during retinal development and for functional maintenance^2,3^. Substitution mutations within the coding region for CRX homeodomain are generally associated with autosomal dominant LCA and CoRD. For example, several studies have evaluated knock-in mouse models harboring three of these mutations, namely *CRX^E80A^*, *CRX^K88N^*, and *CRX^R90W^*. These mutations can alter the DNA-binding capability of mutant CRX proteins, causing distinct pathogenic mechanisms^4-6^. p.E80A binds to the same sites as WT CRX protein but enhances transactivation activities on targets genes such as rhodopsin (*RHO*) at neonatal period^4^. However, p.E80A fails to promote sufficient expression of these genes for subsequent photoreceptor differentiation, with cones being more severely affected^4,6^. p.K88N shows altered DNA-binding specificity, resulting in binding at ectopic sites and perturbation of photoreceptor gene expression during early development^4^. p.R90W exhausts the DNA-binding capability and compromises transcriptional regulation for photoreceptor differentiation. Thus, *CRX^E80A^* and *CRX^K88N^* mutations show antimorphic effects, causing dominant CoRD in *CRX^E80A/+^* patients and LCA in *CRX^K88N/+^* patients^7,8^. The hypomorph *CRX^R90W^*mutation is associated with autosomal recessive LCA^9^. A detailed genotype-phenotype analysis is needed to determine the progression of photoreceptor degeneration and critical windows for potential therapeutic interventions.

Although molecular analysis has gained significant knowledge on how disease mutations disrupt CRX-regulatory activity, the resulting photoreceptor biology and retinal development has not been comprehensively documented in these models, particularly for the detailed course of photoreceptor defects. Defective photoreceptor localizations in many IRDs extend to secondary morphological abnormalities in inner neurons, retinal glia and vasculature^10-14^. This study characterizes the spatiotemporal progression of aberrant retinal morphology in selected mouse models of *CRX* homeodomain mutations. A key observation is that cone differentiation is largely compromised in *Crx^E80A^* and *Crx^K88N^* mutant retinae at young ages. Moreover, *Crx^E80A^* and *Crx^K88N^*mutant retinae develop structural undulations, whorls or folds within neuronal layers, adversely impacting the cellular organization of photoreceptors as well as localizations of inner neurons and retinal vasculature. The analysis of progressive photoreceptor pathologies and secondary morphological defects in these disease models not only provides new insights into IRD pathogenesis but also serves as a paradigm for how misregulated photoreceptor gene expression can disrupt the development of retinal structure and function.

## RESULTS

### Aberrant retinal morphology and development of retinal rosettes

H&E staining of retinal sections is used to evaluate retinal morphology and photoreceptor structure. In the wildtype (*WT*) mouse retina (Figure 1A), H&E staining showed that retinal lamination is established upon the completion of cell proliferation around postnatal day 10 (P10)^15^. The elaboration of outer segments (OS) becomes discernible following eye opening around P14. Synaptic integration and phototransduction is established by approximately P21^16-18^, after which the retinal morphology is preserved through adulthood (after 1 month-old (MO)). In contrast, structural alterations were found in every model of mutant retinae (Figure 1B-E), including focal undulations (Figure 1B), multifocal whorls (Figure 1C, D) and afocal folds (Figure 1E). In the following context, these structural alterations were named as retinal rosettes for simplicity. The *Crx^E80A/+^* retinae (i.e. *E80A/+*, Figure 1B) displayed a developmental trajectory comparable to *WT* controls. However, the wavy architecture emerged at apical outer nuclear layer (ONL) around P14, forming definitive retinal rosettes in adulthood. The *Crx^E80A/A^* retinae (i.e. *E80A/A*, Figure 1C) developed structural turbulence at basal ONL at P10, leading to retinal rosettes with more immense undulations as compared to those in *E80A/+* retinae. Similarly, retinal rosettes were detected at ONL after P10 in the *Crx^K88N/+^* retinae (i.e. *K88N/+*, Figure 1D). Retinal rosettes were pronounced in the *Crx^K88N/N^* retinae (i.e. *K88N/N*, Figure 1E) before P10. Overall, retinal rosettes formed during photoreceptor differentiation, persisted throughout adulthood, and exhibited irregular morphologies and variable sizes. Furthermore, ONL and inner nuclear layer (INL) remained distinct and well-separated in *E80A/+*, *E80A/A* and *K88N/+* retinae, while ONL and INL became assimilated in *K88N/N* retinae from 1 month-old (MO) onward. These morphological features collectively suggested that retinal lamination was initiated during early development in mutant retinae but extensively disrupted by the formation of retinal rosettes. In addition, the structure of OS was missing in *E80A/A*, *K88N/+*, and *K88N/N* retinae, implying the profound defect in photosensitivity and phototransduction.

**Figure 1.**
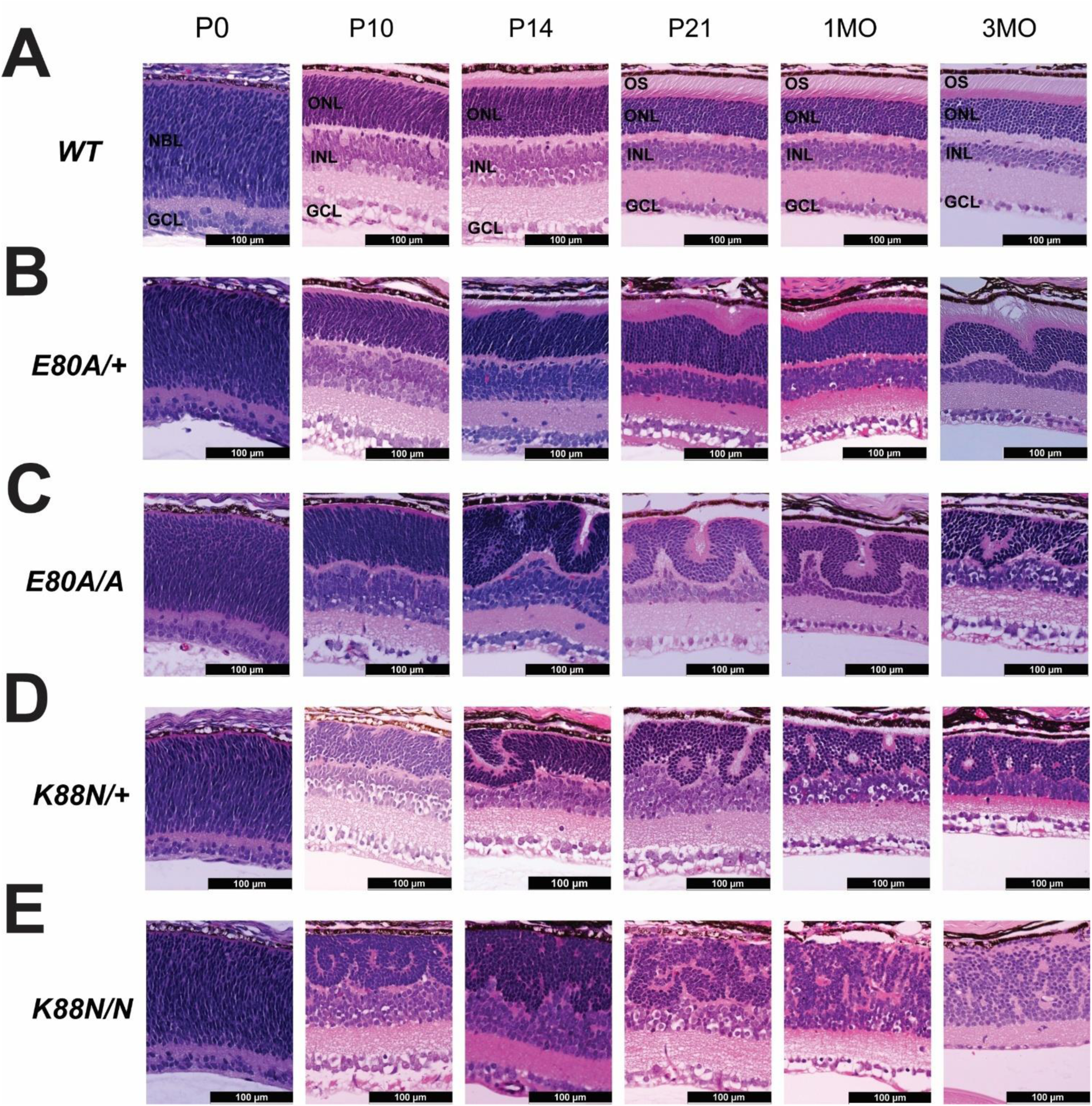
Retinal morphology of *Crx^E80A^* and *Crx^K88N^* mutants. Hematoxylin and eosin (H&E) cross-section staining of (A) wildtype, (B) *Crx^E80A/+^*, (C) *Crx^E80A/A^*, (D) *Crx^K88N/+^*, and (E) *Crx^K88N/N^*retinae at postnatal day 0 (P0), P10, P14, P21, 1 month-old (MO) and 3MO. NBL: neuroblast layer. GCL: ganglion cell layer. ONL: outer nuclear layer. INL: inner nuclear layer. OS: outer segment. Scale bar = 100µm.

### Impaired photoreceptor differentiation and visual function

Photoreceptor morphology, abundance, and localization was examined to determine the condition of photoreceptor differentiation in *Crx^E80A^*and *Crx^K88N^* mutant retinae. Since molecular analysis suggested that cone differentiation was more severely disrupted than rod differentiation, phenotypic characterizations of cone differentiation were reported at first. Retinoid X receptor gamma (RXRG) is expressed in fated cones of neonatal mouse retinae^19,20^. The immunohistochemistry (IHC) staining of RXRG detected cones at apical regions of *WT* and mutant retinae at P0 (Figure 2A). Quantification revealed comparable cone numbers among all genotypes (Supplemental Figure 1A), indicating that cone genesis was unaffected by *Crx^E80A^* and *Crx^K88N^* mutations. However, at P2, RXRG+ cones were scarcely detected in *E80A/A*, *K88N/+*, and *K88N/N* retinae, and only a few were observed in *E80A/+* retinae (Figure 2B, Supplemental Figure 1B). At P10, when cell proliferation concludes in the retina, IHC staining of the cone-marker arrestin 3 (i.e. cone arrestin, ARR3) was detectable only in *WT* and *E80A/+* retinae (Figure 2C). Cell bodies of ARR3+ cones were localized to apical ONL of *WT* retinae but positioned within the middle ONL of *E80A/+* retinae. Similar cone phenotypes were observed in *WT* and mutant retinae at P14 (Figure 2D, Supplemental Figure 2A). Notably, *E80A/+* retinae displayed a complete loss of ARR3+ cones by P21, prior to the completion of photoreceptor differentiation in *WT* retinae. Altogether, this data suggested that p.E80A and p.K88N do not disrupt cone fate specification or the cone precursor pool, instead impair early-stage cone differentiation. Despite their distinct DNA-binding specificities of mutant proteins, *Crx^E80A^* and *Crx^K88N^*mutations both lead to a rapid loss of cone populations shortly after birth in *E80A/A* and *K88N/N* mutant retinae. Even though *K88N/+* retinae carried one *WT Crx* allele, the ectopic DNA binding of p.K88N impaired early-stage cone differentiation during the critical window of P0 to P2. *E80A/+* mutant retinae showed a progressive loss of cones during retinal development (Supplemental Figure 2B), supporting the molecular evidence that p.E80A fails to establish the proper chromatin landscape for sufficient expression of cone phototransduction genes for later differentiation^6^.

**Figure 2.**
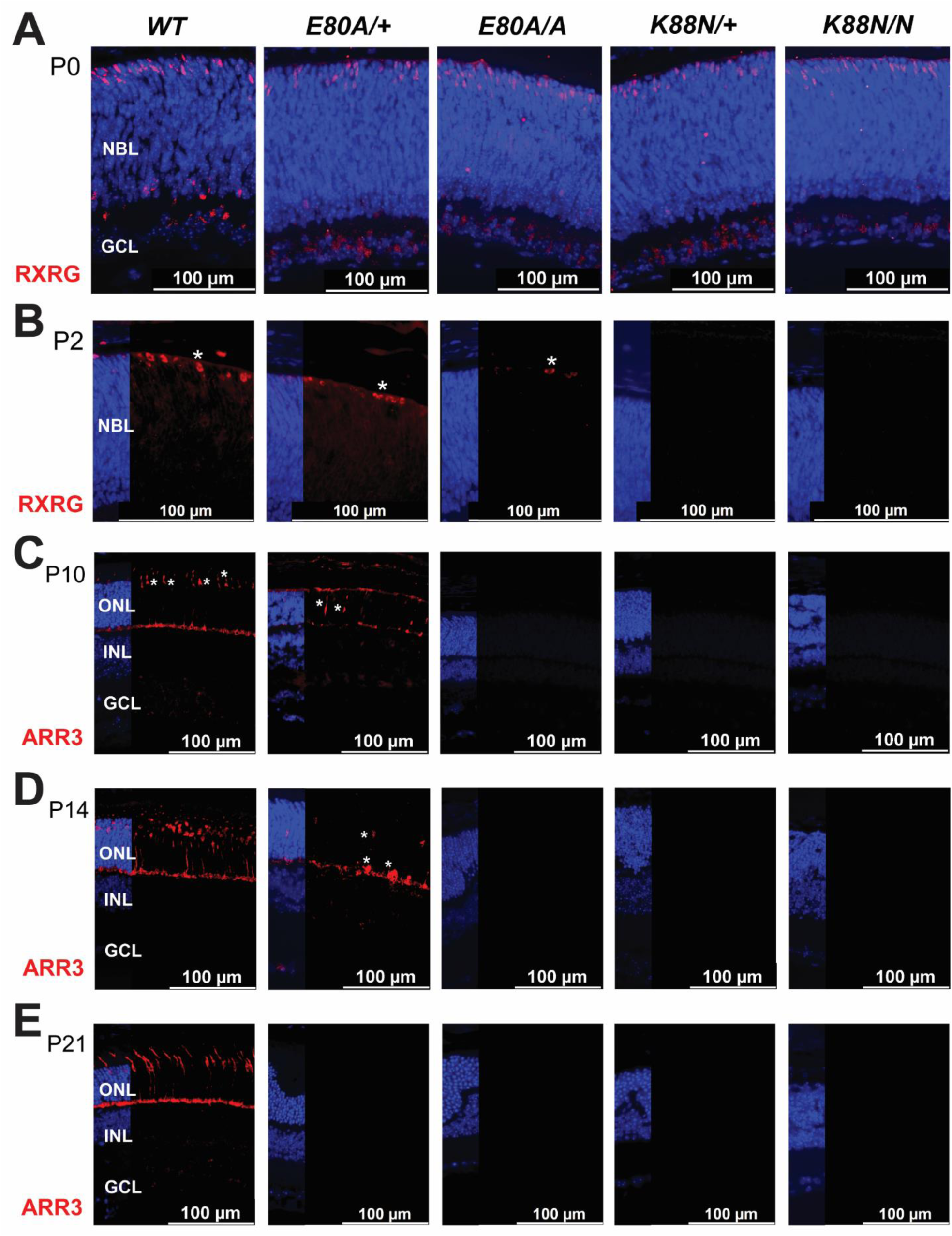
Defective cone differentiation in *Crx^E80A^* and *Crx^K88N^* mutants. Immunohistochemistry (IHC) staining of RXRG (in red) in (A) P0, (B) P2 *WT* and mutant retinae. IHC staining of ARR3 (in red) in (C) P10, (D) P14, (E) P21 *WT* and mutant retinae. Nuclei are stained by DAPI (in blue). Scale bar = 100µm. Asterisks indicate examples of labelled cones.

In contrast to cones, diseased rods in mutant retinae underwent a different phenotypic trajectory. IHC staining of rhodopsin (RHO) was sporadically detected in *WT* retina but found significantly enriched in the *E80A/+* and *E80A/A* retinae at P3 (Figure 3A), suggesting that p.E80A caused premature rod differentiation relative to p.WT and p.K88N^4^. At P10, RHO expression was localized in ONL of *WT*, *E80A/+*, *E80A/A*, and *K88N/+* retinae, but became absent in *K88N/N* retinae (Figure 3B). At P21, RHO expression was found within OS of *WT* and *E80A/+* retinae, while RHO expression was localized within the disorganized ONL of *E80A/A* and *K88N/+* retinae (Figure 3C). Furthermore, a *Nrl* promoter-driven cytoplasmic GFP reporter (*pNrl-eGFP*, a rod lineage marker) was detectable in ONL of *E80A/+*, *E80A/A* and *K88N/+* retinae at P21; GFP expression was lost in *K88N/N* retinae, suggesting that the rod lineage commitment was disturbed with minimal activity of *Nrl* promoter (Figure 3D). In summary, cone differentiation was more severely affected than rod differentiation in mutant retinae, besides *K88N/N* retinae showing no sign of rod differentiation. RHO expression appeared precociously in neonatal *E80A/+* and *E80A/A* retinae; RHO expression was later trafficked to OS of *E80A/+* retinae but retained in ONL of *E80A/A* retinae that failed to develop OS. *K88N/+* retinae also had RHO expression mislocalized in ONL.

**Figure 3.**
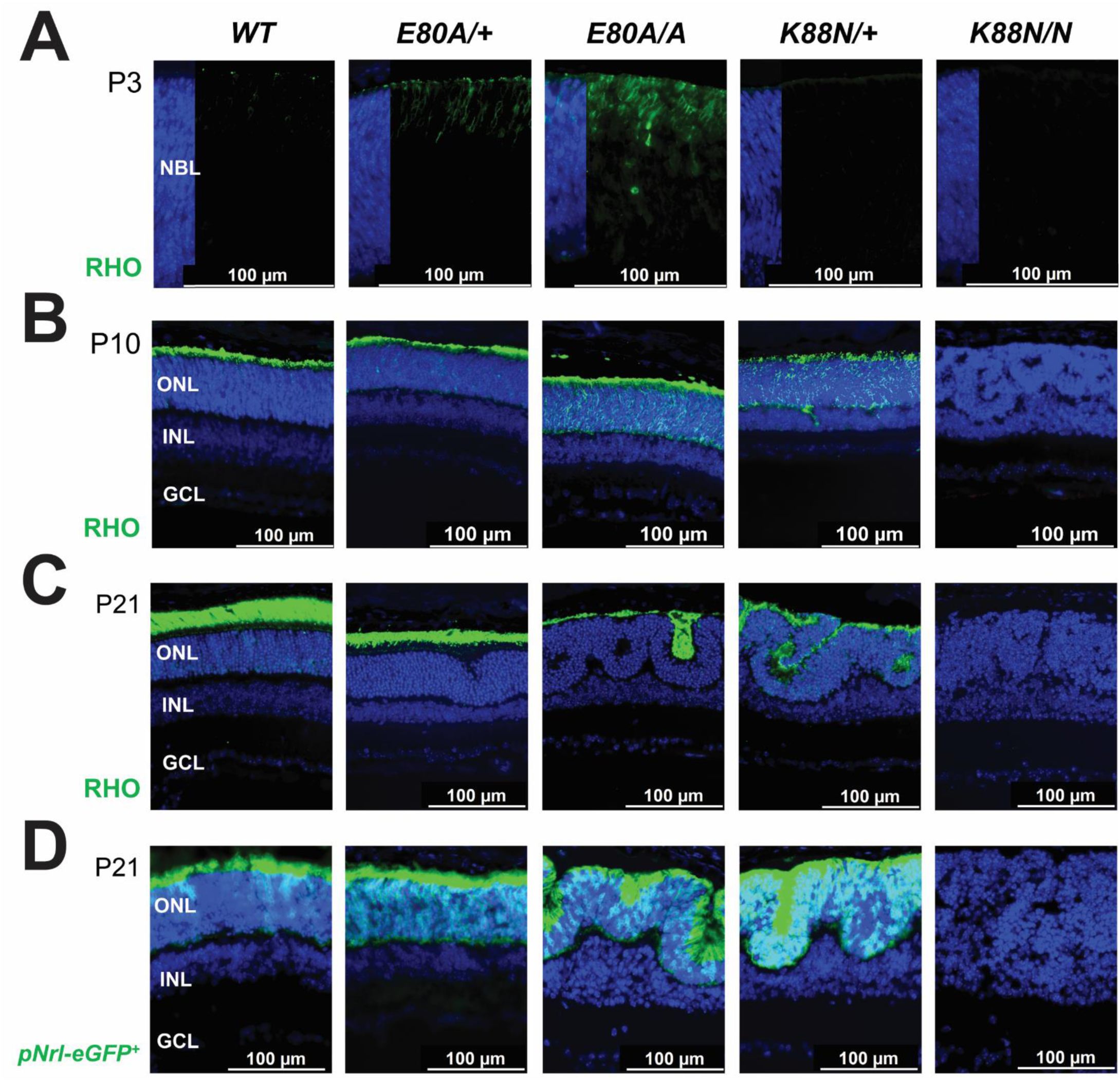
Incomplete rod differentiation in *Crx^E80A^* and *Crx^K88N^* mutants. IHC staining of RHO (in green) in (A) P3, (B) P10, (C) P21 *WT* and mutant retinae. (D) Green cells labelled by *pNrl-eGFP* in P21 *WT* and mutant retinae. Nuclei are stained by DAPI (in blue). Scale bar = 100µm.

Apoptosis analysis with cleaved caspase 3 (CC3) staining at P2, P10, P14 and P21 showed a significant increase only in the *K88N/N* retina at P21, but strikingly, found non-significant difference in other comparisons (Supplemental Figure 3A-D, F). These data suggested that initial photoreceptor loss in developing mutant retinae was implausibly driven by cell death. Apoptosis analysis at 3MO reported increased cell death in mutant retinae (Supplemental Figure 3E, F). Notably, apoptotic cells could be occasionally found in apical ONL bearing retinal rosettes.

Electroretinography (ERG) was performed to determine if disrupted photoreceptor differentiation and retinal structure impaired visual functions in adult mice. Previous studies reported that *E80A/A*, *K88N/+*, and *K88N/N* mice lacked ERG responses at 1MO^4^. As expected, ERG response remained undetectable in these mutants at 3MO (Supplemental Figure 4A-C). Rod-driven responses (dark-adapted A and B waves) were reduced but detectable in 1MO *E80A/+* mice, but cone-driven responses (light-adapted B waves) were however missing^4^. Although H&E staining revealed the presence of OS structure, 3MO *E80A/+* retinae showed greatly reduced to near-complete loss of rod-driven responses as compared to *WT* controls (Supplemental Figure 4A, B). Cone-driven responses remained nought in 3MO *E80A/+* mice (Supplemental Figure 4C). A longitudinal comparison of rod-driven ERG responses in *E80A/+* samples reported a progressive loss of rod function between 1MO and 3MO (Supplemental Figure 4D). IHC staining was employed to determine cellular mechanisms underlying the reduced visual function in 3MO *E80A/+* retinae. RHO expression was localized to OS of 1MO and 3MO *E80A/+* retinae (Supplemental Figure 5A), including within regions containing retinal rosettes. However, GNAT1, a key transducing protein of the rod phototransduction cascade^17,21,22^, was detected in OS of 1MO *E80A/+* retinae but significantly decreased by 3MO (Supplemental Figure 5B). VGLUT1 expression is usually detected in the presynaptic sites^23,24^. The protrusion of retinal rosettes in *E80A/+* retinae impeded VGLUT1 expression (Supplemental Figure 5C, white arrows), in contrast to adjacent regions without retinal rosettes (Supplemental Figure 5C, yellow arrows). These data suggested that the loss of ERG responses in *E80A/+* mutant retinae likely resulted from the compound defects in phototransduction and synaptic connectivity.

Quantitative PCR (qPCR) analysis provided molecular validation for cellular and functional phenotypes. At neonatal ages, all mutant genotypes showed comparable expression of *Crx* and *Nrl* transcripts to those of *WT* controls (Supplemental Figure 6A), except a moderate downregulation of *Nrl* expression in P5 K88N/N retinae. *Rxrg* expression was comparable between all mutant genotypes and *WT* controls at P0 but became severely reduced in mutant retinae at P3 (Supplemental Figure 6A). During photoreceptor differentiation, *Rho* and *Gnat1* expression underwent a progressive downregulation, dropping to less than 50% of *WT* controls in 1MO *E80A/+* retinae. This transcriptional downregulation mechanistically correlated with the reduced rod-driven ERG responses^4^ (Supplemental Figure 6B). Cone opsin expression was rapidly ablated during photoreceptor differentiation (Supplemental Figure 6B), confirming the degeneration phenotypes and prior sequencing findings^6^.

### Unaffected development of inner neurons

Since photoreceptors closely interact with inner neurons, the next question asked if impaired photoreceptor differentiation caused secondary morphological changes in inner neurons of the *E80A* and *K88N* mutant retinae. The cell abundance and localization of inner neurons was examined by IHC staining at key age points during photoreceptor differentiation. Horizontal cells were labelled by calbindin D28K staining^25^, rod bipolar cells by protein kinase C alpha (PKCα)^26^, amacrine cells by calretinin^27^, retinal ganglion cells by calretinin and RNA-binding protein with multiple splicing (Rbpms)^28^. At P10 prior to the formation of retinal rosettes, inner neurons were correctly organized in mutant retinae (Supplemental Figure 7A-C), and their cell numbers were comparable to those in *WT* controls (Supplemental Figure 7D). Occasionally, mislocalized cells were observed in *K88N* retinae (Supplemental Figure 7B, stars). These data suggested that the developmental programs of inner neurons were minimally affected by impaired photoreceptor differentiation in *E80A* and *K88N* mutant retinae. At P21, the protrusion of retinal rosettes impacted the localizations of horizontal cells and rod bipolar cells in the *E80A/+*, *E80A/A*, and *K88N/+* retinae, although the cell count analysis found non-significant difference between *WT* and mutant retinae (*E80A/+*, *E80A/A*, and *K88N/+* in Figure 4A, B, E). Interestingly, fewer labelled cells were found within regions in contact with protruded ONL (Figure 4A, B, white arrows) as compared to adjacent regions without retinal rosettes in the *E80A/A* and *K88N/+* retinae (Figure 4A, B, yellow arrows). Horizontal cells and rod bipolar cells were present in *K88N/N* retinae, although their cell numbers were reduced as compared to those in *WT* controls (*K88N/N* in Figure 4A, B, E). Mislocalized cells were again evident in P21 *K88N/N* retinae (Figure 4A, B, stars). In contrast, the cell abundance and localization of amacrine cells and ganglions cells were unaltered in mutant retinae (Figure 4C, D, E). In conclusion, the presence of retinal rosettes induced morphological alterations in neighboring horizontal cells and rod bipolar cells in developing *E80A/+*, *E80A/A*, and *K88N/+* retinae; amacrine cells and ganglions cells were relatively distant from retinal rosettes, thus remaining unaffected. *K88N/N* retinae showed a reduction in cell numbers of horizontal cells and rod bipolar cells, possibly due to the worse structural disorganizations.

**Figure 4.**
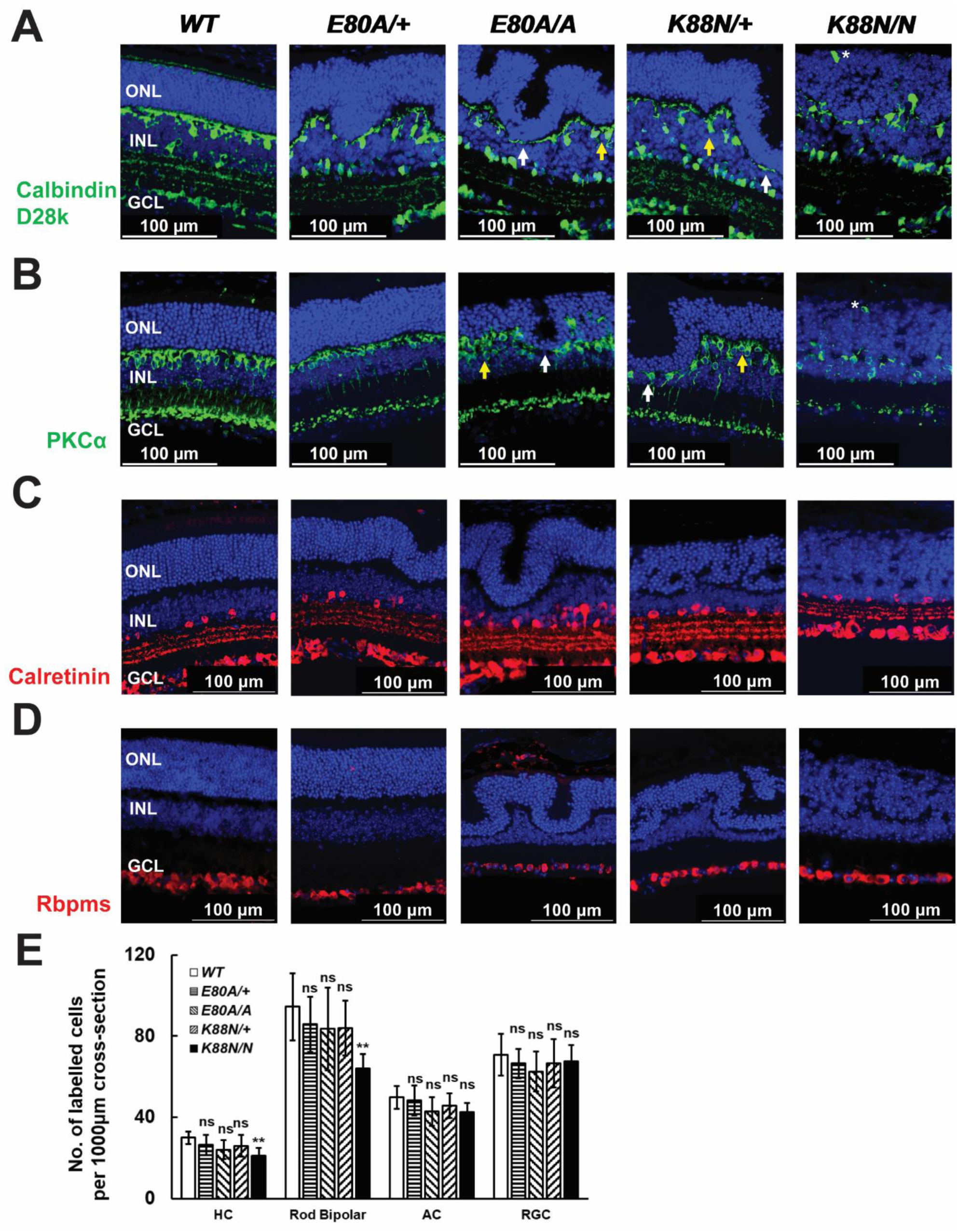
Inner neurons in *Crx^E80A^*and *Crx^K88N^* mutants. IHC staining of (A) calbindin D28K (in green), (B) PKCα (in green), (C) calretinin (in red), (D) rbpms (in red) in P21 *WT* and mutant retinae. White arrows indicate regions lacking labelled cells. Yellow arrows indicate clusters of labelled cells. Nuclei are stained by DAPI (in blue). Scale bar = 100µm. (E) Cell counts for different retinal cell types between 500µm and 1500µm cross-sections of P21 *WT* and mutant retinae. Error bars represent SD on mean values (n=4). Statistical analysis by pairwise t-test happens between *WT* and each mutant group. HC: horizontal cells. AC: amacrine cells. RGC: retinal ganglion cells. Asterisks (**) denote p≤0.05, ns means not significant.

### Moderate activation of retinal glia

Reactive gliosis can be triggered by photoreceptor injury during retinal development^14^. Müller glia and microglia were analyzed to assess the potential of glial activation in *Crx^E80A^* and *Crx^K88N^*mutant retinae at key age points during photoreceptor differentiation using immunohistochemistry. Glutamine synthetase (GS) is an enzyme specifically localized in Müller glia, and its immunoreactivity covers the radial morphology that traverses from outer limiting membrane (OLM) to inner limiting membrane^29^. At P21, GS immunoreactivity was readily detected in mutant retinae (Figure 5A), implying the presence of Müller glia. In particular, GS immunoreactivity in *E80A/+*, *E80A/A*, *K88N/+* retinae extended apically and basally to span the retinal width, following the shape of undulations. However, GS immunoreactivity appeared misoriented and did not display the radial extension in *K88N/N* retinae. Cell bodies of Müller glia were labelled by P27^Kip1^ staining^30^. Müller glia were mislocalized in *E80A/A* and *K88N/+* retinae (*E80A/A* and *K88N/+* in Supplemental Figure 8A), mirroring the disarray of horizontal cells and rod bipolar cells, i.e. fewer Müller glia within regions in contact with protruded ONL (Supplemental Figure 8A, white arrows) but more at adjacent regions without retinal rosettes (Supplemental Figure 8A, yellow arrows). Cell bodies of Müller glia appeared clustered in *K88N/N* retinae (*K88N/N* in Supplemental Figure 8A), and cell count analysis found a significant decrease in cell numbers of Müller glia in *K88N/N* retinae (Supplemental Figure 8B). In response to acute injury or stress, Müller glia become reactive and accumulate glial fibrillary acid protein (GFAP)^14^. At P21, GFAP expression was found at nerve fiber layer in all mutants, likely attributed to astrocytes. *K88N/N* retinae displayed uneven upregulation of GFAP expression, whereas GFAP expression was weakly detectable in other mutant retinae (Figure 5B). Overall, the moderate GFAP expression indicated a limited reactive gliosis in mutant retinae.

**Figure 5.**
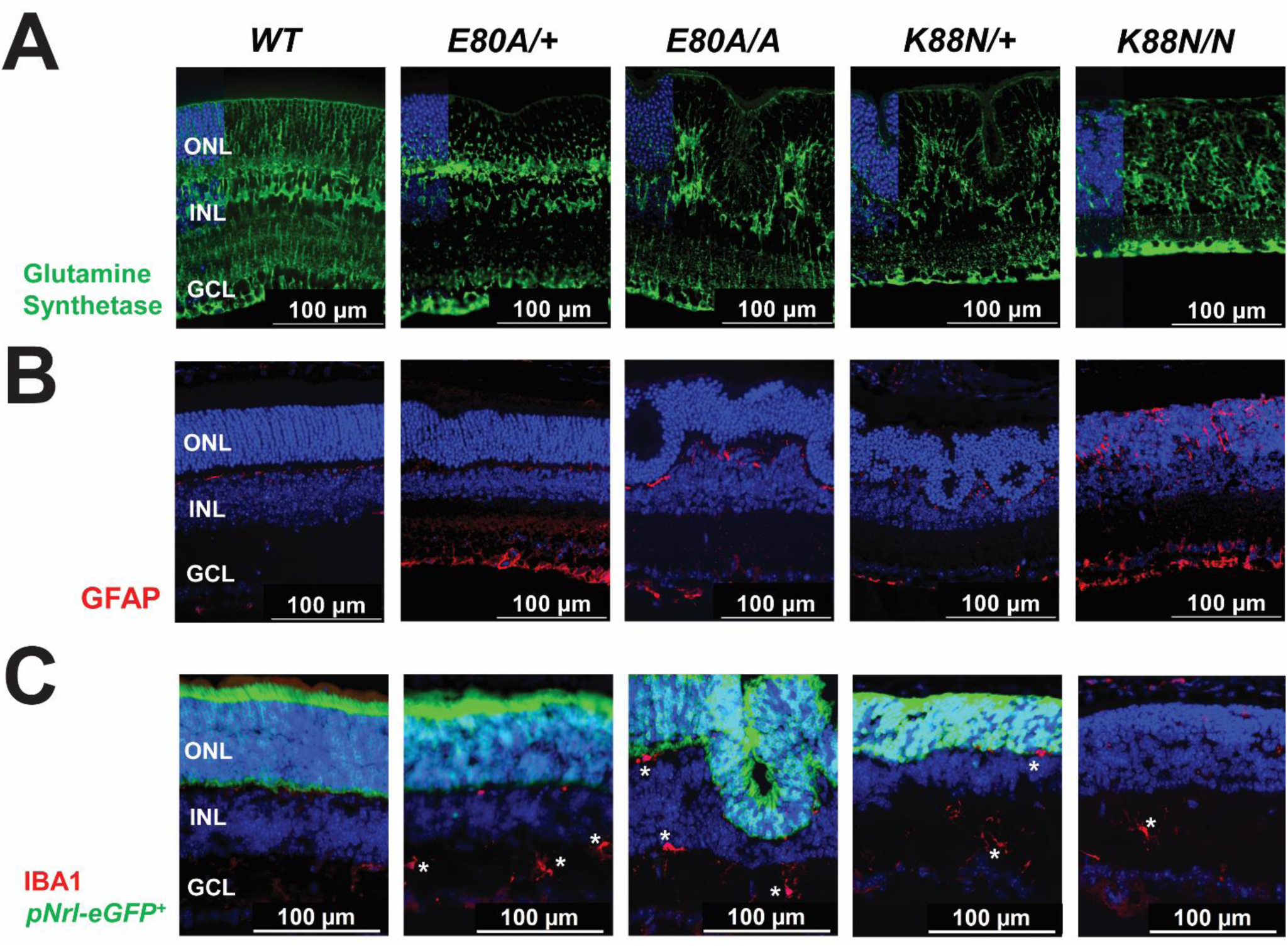
Retinal glia in *Crx^E80A^*and *Crx^K88N^* mutants. IHC staining of (A) glutamine synthetase (in green), (B) GFAP (in red), (C) IBA1 (in red) in P21 *WT* and mutant retinae. Green cells in (C) are labelled by *pNrl-eGFP*. Asterisks in (C) indicate examples of IBA1+ cells. Nuclei are stained by DAPI (in blue). Scale bar = 100µm.

Activated microglia phagocytose damaged, dying, misconnected, or mislocalized retinal neurons^13^. Ionized calcium binding adaptor molecule 1 (IBA1) functions as a microglia and macrophage-specific calcium binding protein and becomes a specific marker for microglia in the retina^31,32^. *pNrl-eGFP* was employed to accurately identify ONL (especially for regions of retinal rosettes) from INL in *WT*, *E80A/+*, *E80A/A*, and *K88N/+* retinae. Although the mutant retinae contained numerous mislocalized cells within retinal rosettes, microglia were detected exclusively in outer plexiform layer (OPL), INL, and inner plexiform layer (IPL) (Figure 5C, stars). In addition, P2Y12 receptor (P2Y12R) staining confirmed that microglia were localized outside retinal rosettes (Supplemental Figure 9A, stars). Interestingly, CD68 staining found infiltrated cells, likely macrophages, at the outer surface of retinal rosettes in *E80A/A* and *K88N/+* retinae and within neuronal layer of the *K88N/N* retinae (Supplemental Figure 9B, stars), suggesting that phagocytic macrophages may be involved in the structural formation of retinal rosettes.

### Abnormal neurovascular networks

In a retina, superficial vascular plexus (SVP), intermediate vascular plexus (IVP), and deep vascular plexus (DVP) form three layers of the retinal capillary network. SVP contains a network of blood vessels located in the retinal ganglion cell layer and nerve fiber layer. IVP is located within IPL, branching out to connect SVP and DVP. DVP resides in OPL and contacts with photoreceptors. IVP and DVP develop concurrently with photoreceptor differentiation^33,34^. To this end, vascular development was analyzed in *Crx^E80A^* and *Crx^K88N^* mutant retinae. Conjugated lectin staining revealed the vascular endothelium at key age points during photoreceptor differentiation. *pNrl-eGFP* reporter was employed to distinguish ONL from INL in the *WT*, *E80A/+*, *E80A/A*, and *K88N/+* retinae. At P10, vessels at SVP and DVP were correctly localized in *E80A/+*, *E80A/A*, *K88N/+* retinae, as compared to those in *WT* controls (Figure 6A). In contrast, the retinal vasculature was profoundly disorganized in the *K88N/N* retina. Vascular elements were ectopically localized within retinal rosettes; vessels traversed the neuronal layer to reach OLM (Figure 6A, yellow arrow). Interestingly at P21, vessels at DVP remodeled in parallel with the protrusion of retinal rosettes in the *E80A/+*, *E80A/A*, and *K88N/+* retinae (Figure 6B, white arrows). Vessels in *K88N/N* retinae remained in a disorganized clutter at P21, with aberrant vessels penetrating the entire neuronal layer (Figure 6B, yellow arrow). The wholemount image at OLM identified ectopic vascular networks in *K88N/N* retinae (Supplemental Figure 10, yellow arrows). The 3D reconstruction of vasculature between INL and ONL confirmed the presence of DVP and IVP in *WT*, *E80A/+*, *E80A/A*, and *K88N/+* retinae (Figure 7A-D). Rather than forming flat plexuses, vessels aberrantly wrapped around the protrusion of retinal rosettes in mutant retinae (Figure 7B’-D’). Besides vessels at regions of retinal rosettes, the structural components of vasculature in *E80A/+*, *E80A/A*, and *K88N/+* mutant retinae resembled those in *WT* retinae. However, neither DVP nor IVP could be found in the vascular landscape of *K88N/N* retinae. Many vessels instead grew along the apical-basal axis (Figure 7E, E’) in *K88N/N* retinae. Vascular endothelial growth factor receptor 2 (VEGFR2) is known to regulate endothelial migration and proliferation^35,36^. At P14, VEGFR2 expression was localized to regions with evolving retinal rosettes in *E80A/A*, *K88N/+*, and *K88N/N* retinae (Supplemental Figure 11, white arrows), indicating that VEGFR2-mediated signaling may be involved in coupled pathologies of DVP disorganization and the formation of retinal rosettes. Collectively, these data support a close 3-way non-autonomous interaction between photoreceptor differentiation, the formation of retinal rosettes, and the aberrant organization of retinal vasculature in mutant retinae.

**Figure 6.**
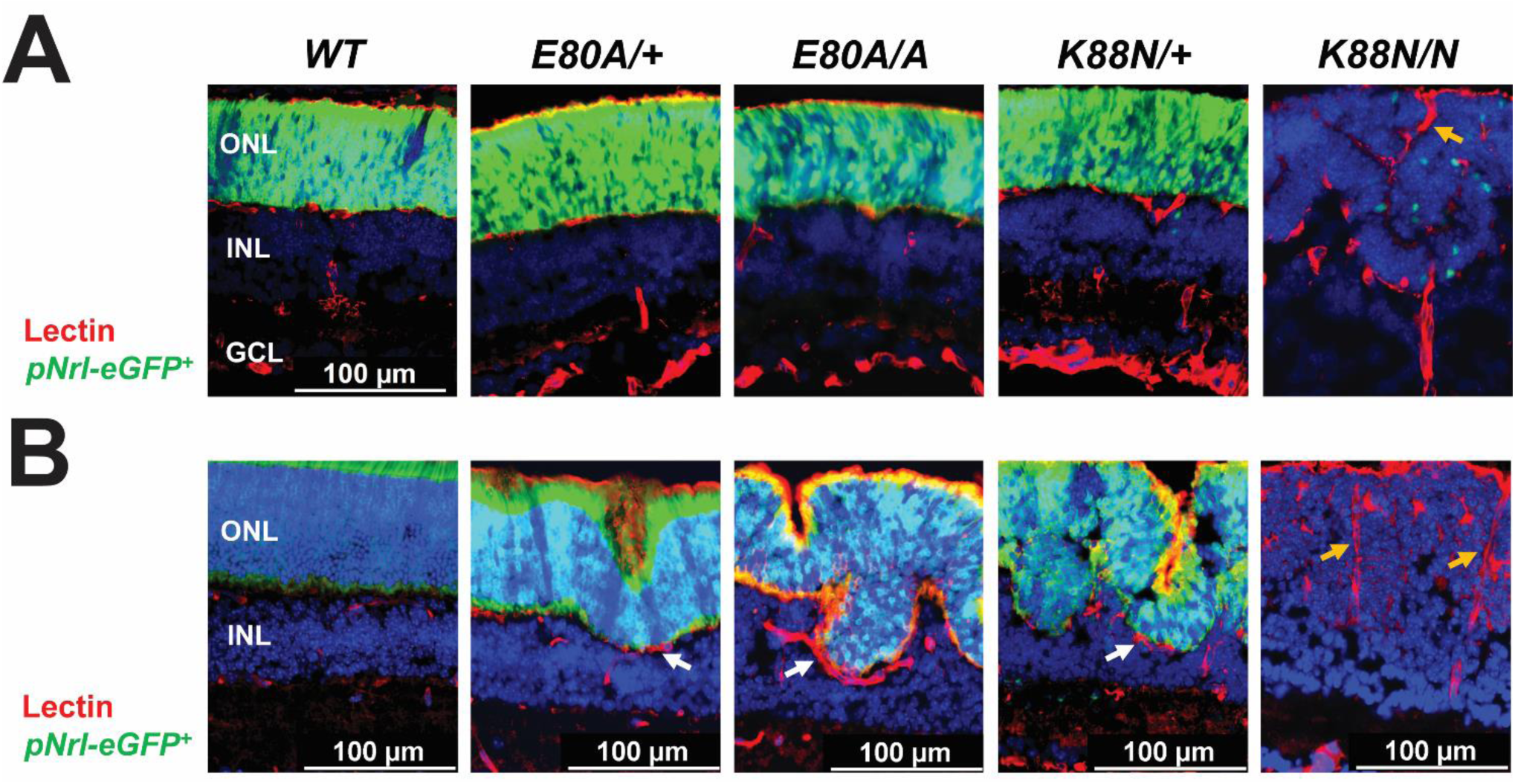
Vasculature in *Crx^E80A^*and *Crx^K88N^* mutants. IHC staining of conjugated lectin (in red) in (A) P10, (B) P21 *WT* and mutant retinae. Green cells are labelled by *pNrl-eGFP*. White arrows point to vessels wrapping around retinal rosettes. Yellow arrows indicate vessels running through neuronal layers of *Crx^K88N/N^* retinae. Nuclei are stained by DAPI (in blue). Scale bar = 100µm.

**Figure 7.**
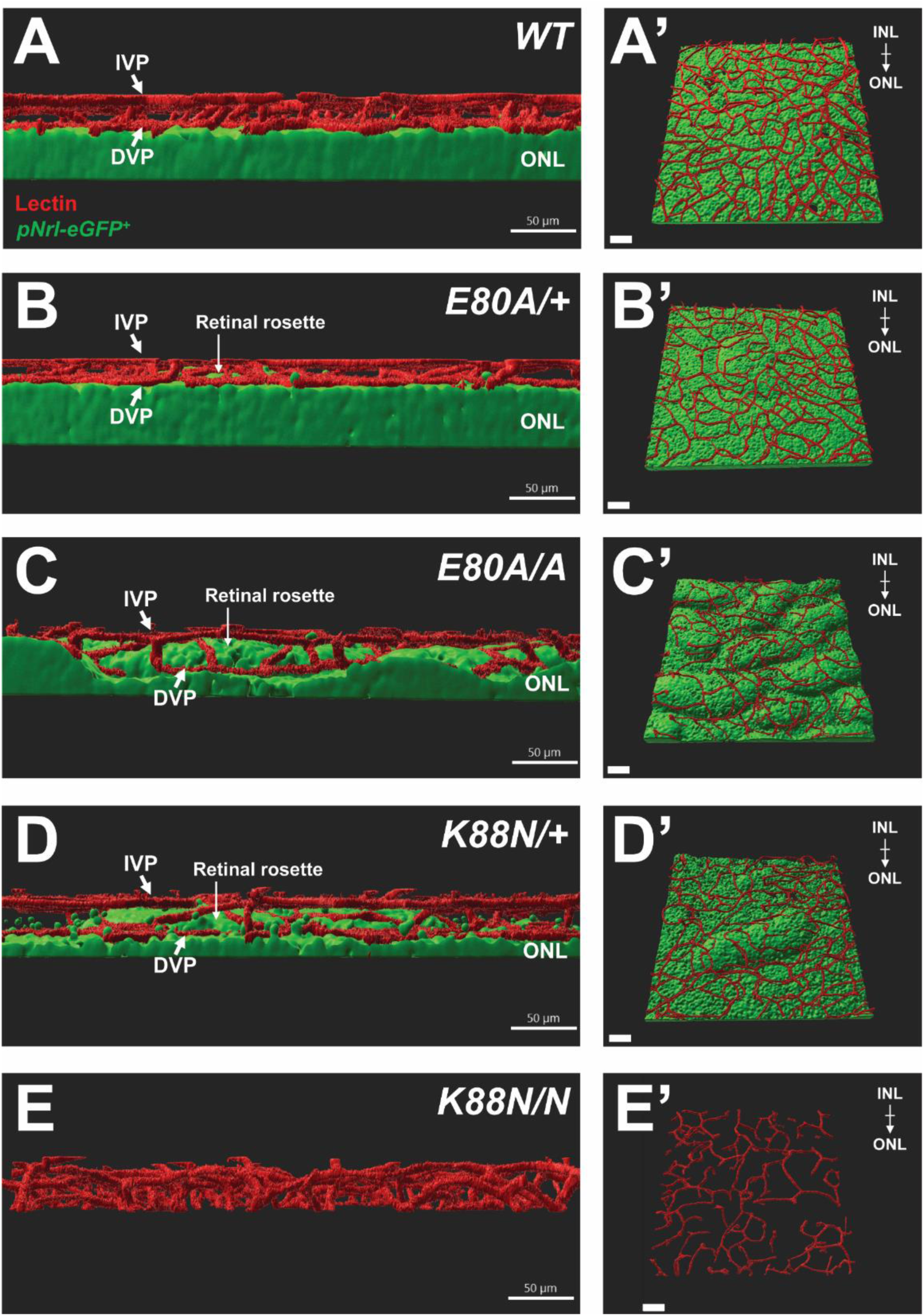
Vasculature landscape in *Crx^E80A^* and *Crx^K88N^* mutants. (A-E) Intermediate vascular plexus (IVP) and deep vascular plexus (DVP) in cross-sections of P21 *WT* and mutant retinae. ONLs in (A-D) are labelled green. Retinal rosettes are pointed in (B-D). (A’-E’) Top view oriented from INL towards ONL.

### Unaffected retinal lamination and vasculature in CRX-loss-of-function mutants

The next question asked if the formation of retinal rosettes and aberrant vasculature was specific cellular signatures of the specific *Crx* homeodomain mutations *Crx^E80A^* and *Crx^K88N^*, or a common consequence of incomplete photoreceptor differentiation. The loss-of-function *CRX^R90W^* mutation produced a mutant protein devoid of DNA-binding activity, similar to the case of *Crx^-/-^*^4,5^. Photoreceptor differentiation was absent in *Crx^R90W/W^* (i.e. *R90W/W*) and *Crx^-/-^*retinae (Supplemental Figure 12A). These retinae did not form retinal rosettes. At P21, horizontal cells (Supplemental Figure 12B), rod bipolar cells (Supplemental Figure 12C), amacrine cells, retinal ganglions cells (Supplemental Figure 12D), and Müller glia (Supplemental Figure 12E) were present in *R90W/W* and *Crx^-/-^* retinae, suggesting the development programs of these cell types were unaffected. GFAP was moderately activated in *R90W/W* and *Crx^-/-^* retinae (Supplemental Figure 12F), and some IBA+ cells were found at OPL, INL and IPL in mutant retinae (Supplemental Figure 12G). Lectin staining detected three plexuses in *R90W/W* and *Crx^-/-^* retinae (Supplemental Figure 12H). Therefore, these loss-of-function mutations result in ablated photoreceptor differentiation but do not disrupt the developmental programs or organizations of inner neurons, glia, and retinal vasculature.

## DISCUSIONS

*CRX^E80A^*, *CRX^K88N^*, and *CRX^R90W^*mutations cause distinct effects on photoreceptor differentiation. In *E80A/+* retinae, cone genesis appears unaffected, but the subsequent differentiation is progressively diminished. In contrast, rod differentiation initiates earlier in *E80A/+* retinae, and subsequent elaboration of the OS structure develops normally. However, rods in *E80A/+* retinae fail to maintain functionality in adulthood.

Cone differentiation is absent in *K88N/+* retinae shortly after birth, and rod differentiation fails to develop functional rods. These phenotypes in mouse models recapitulate the clinical manifestations, i.e. dominant cone-rod dystrophy in *CRX^E80A/+^* patients and dominant Leber congenital amaurosis in *CRX^K88N/+^* patients^7,8^. *E80A/A* retinae show signs of incomplete rod differentiation but ablated cone differentiation, while *K88N/N* retinae exhibit no photoreceptor differentiation. These phenotypes reveal cell-type specific effects on transcriptional regulation: p.E80A still regulates *Rho* expression but fails to activate cone differentiation; p.K88N represses photoreceptor differentiation in general.

Photoreceptor pathologies in *Crx^E80A^* and *Crx^K88N^*mutant retinae suggest important features of photoreceptor differentiation. Firstly, cone differentiation has a critical time window shortly after the event of fate choice to establish and maintain cone identity. Secondly, cone differentiation requires a fine-tuned and precise transcriptional program, for which neither hyperactive (p.E80A) nor ectopic (p.K88N) activity can be tolerated^6^. Thirdly, rod differentiation can be defined by three stages, namely, the establishment of cell identity, OS elaboration, and phototransduction. Results of RHO staining and *pNrl-eGFP* fluorescence suggest that rod identity can be retained even under hyperactive transcriptional activity. However, OS elaboration needs more stringent control on transcriptional activity: at least a copy of WT allele has to be present. Phototransduction demands an orchestra of various components in cell body and OS, which can only be maintained by sufficient activity of p.WT CRX protein. Future work will be devoted to further investigation of affected components of OS, phototransduction and synaptic connectivity due to these mutations. Fourthly, photoreceptor differentiation is completely lost in *K88N/N* and *R90W/W* retinae. However, *R90W/W* retinae undergo a rapid ONL degeneration to a complete loss before 3MO^5^; cells survive much longer in the assimilated layer of *K88N/N* retinae. This difference in cell survival suggests that affected cells may adopt distinct cellular states or responses to stress. Lastly, since heterozygous mutants of each genotype show less severe phenotypes than homozygous ones, *CRX* augmentation may represent a potential treatment strategy^37^. Ectopic expression of *WT* CRX may provide the essential transcriptional activity to drive photoreceptor differentiation in *CRX^E80A^* and *CRX^K88N^* mutant retinae. Due to altered binding activities of mutant CRX proteins, optimal efficacy may require a dual strategy of combining *CRX* augmentation with genetic suppression of the mutant alleles to counteract the antimorphic effects. All in all, this study provides comprehensive phenotypic evidence to established molecular mechanisms on *CRX^E80A^*, *CRX^K88N^* mutations^4,6^ and identifies critical windows for potential therapeutic interventions.

Phenotypes of retinal rosettes in *CRX^E80A^* and *CRX^K88N^*mutant retinae are pronounced. In *Nrl^-/-^* and *Nr2e3^rd7/rd7^*mice, retinal rosettes are formed in cone-dominant ONL or arise from disrupted rod differentiation^38,39^. Interestingly, *E80A/A* and *K88N/+* retinae develop retinal rosettes in rod-dominant ONL. Regardless of ONL cellular identity, *Nrl^-/-^*, *Nr2e3^rd7/rd7^*, *Crx^E80A/A^*, *Crx^K88N/+^* mice develop retinal rosettes at approximately the same age. A question thus arises: why does impaired photoreceptor differentiation in transcription factor mutants lead to the formation of retinal rosettes? Although the detailed mechanisms remain to be elucidated, this study provides cellular insights to further address this question. Firstly, OS elaboration in *E80A/+* retinae concurrently occurs with ONL undulations around P14, while a few cones are still detectable at this age. Hence, at least in the cases of *CRX^E80A^* and *CRX^K88N^* mutant retinae, a cone-dominant or rod-dominant composition is not a specific characteristic to the formation of retinal rosettes. Secondly, *K88N/N* retinae exhibit a complete loss of photoreceptor identity and differentiation, developing a more severe form of retinal rosettes before P10. In contrast, CRX-loss-of-function *R90W/W* and *Crx^-/-^* retinae do not develop retinal rosettes, even though their ONL cells retain immature photoreceptor identity. These findings suggest that the formation of retinal rosettes cannot be explained solely by the conditions of photoreceptor identity or maturation. Therefore, the formation of retinal rosettes is likely associated with misregulated transcription by the mutant p.E80A and p.K88N. It is worth mentioning that this study does not specifically address if perturbed nuclear condensation contributes to the formation of retinal rosettes during photoreceptor differentiation. Based on H&E and DAPI staining, *E80A/+* retinae show signs of nuclear condensation, yet still develop retinal undulations. It is unclear if *E80A/A* and *K88N/+* retinae undergo any form of nuclear condensation, despite showing RHO expression within retinal rosettes. Nuclear condensation appears lost in *K88N/N* retinae. Future studies will comprehensively analyze the nuclear morphology of ONL neurons in these mutants during photoreceptor differentiation.

Phenotypes of inner neurons, retinal glia and vasculature are of particular interest and significance. Firstly, the localizations of Müller glia, horizontal cells and rod bipolar cells are ‘pushed’ aside by the presence of retinal rosettes. Secondly, mislocalized cells within retinal rosettes can escape microglia scavenger during photoreceptor differentiation and induce only moderate GFAP reaction. Lastly, vascular outgrowth around retinal rosettes interacts with nearby neurons via local signaling pathways in *E80A/+*, *E80A/A*, and *K88N/+* retinae. The aberrant vasculature in mutant retinae likely provides nutrients and oxygen to mislocalized cells within retinal rosettes, thereby supporting their survival. Future studies will analyze the molecular mechanisms underlying the remodeling of inner neurons, retinal glia and vasculature in these models and the crosstalk between defective photoreceptors and these cells.

Vessels in *K88N/N* retinae become disorganized and penetrate through the neuronal layers as early as P10. A similar phenotype of vascular penetration can be seen in ONL of *Vldlr^-/-^* retinae during adulthood^40^. To this end, *K88N/N* retinae may serve as a new model to study retinal angiomatous proliferation in absence of photoreceptor differentiation. Retinal choroidal anastomosis is commonly seen in *K88N/N* retinae, which requires further studies to identify the cellular underpinning.

An unanswered question remains if the rosette formation and vascular defects observed in this study are recapitulated in human pathology. It is known that IRDs, especially early-onset ones, cause progressive vision loss in conjunction with significant vascular complications^41,42^. This study identifies a potential cellular link between incomplete photoreceptor differentiation and vascular defects. These findings suggest that early intervention targeting vascular abnormalities should be considered in IRD therapeutic strategies.

In summary, this study reports non-autonomous disease complications that impaired photoreceptor differentiation drives the formation of retinal rosettes, which in turn secondarily disrupts the organization of the retinal vasculature. These findings highlight the necessity of considering vascular pathologies for designing potential therapeutic strategies.

## MATERIALS AND METHODS

### Transgenic Mouse Lines

All mice in this study were on the genetic background of C57BL/6J (the Jackson Laboratory, Stock No: 000664). *CRX^E80A^*, *CRX^K88N^*, and *CRX^R90W^* mutant mice, as well as *pNrl-eGFP* mice were generated for previously published studies^4,5,43^. Both male and female mice were used in experiments.

### Retinal Histology

Eyes were dissected at various ages with the superior/dorsal side marked on the cornea and fixed in 4% paraformaldehyde at 4°C overnight. Tissues were processed with paraffin embedding. Each retinal cross-section was cut by 4 microns on a microtome. Hematoxylin and eosin (H&E) staining was performed to examine retinal histology.

### Immunohistochemistry (IHC) Staining

A summary of cell markers and antibodies involved in this study is detailed in Table 1. IHC staining was performed in paraffin-embedded and cryosections based on the protocol used in the previous study^44^.

**Table 1.** Antibody information.

| Antibody | Product Information | Target | Sampling Age |
| --- | --- | --- | --- |
| RXRG | Santa Cruz Biotechnology, sc-555 | Fated cones | P0 |
| RHO | MilliporeSigma, O4886 | Rods | After P3 |
| Cleaved caspase 3 | Cell Signaling Technology, 9661 | Apoptotic cells | All ages |
| ARR3 | MilliporeSigma, AB15282 | Cones | After P10 |
| GNAT1 | MilliporeSigma, GT40562 | Rod OS | 1MO, 3MO |
| VGLUT1 | MilliporeSigma AB5905 | Presynaptic terminals | 1MO, 3MO |
| Calbindin D28K | MilliporeSigma, C9848 | HCs | P10 |
| Calbindin D28K | MilliporeSigma, ABN2192 | HCs, some ACs & RGCs | P21 |
| PKCα | MilliporeSigma, P4334 | Rod BCs | P10, P21 |
| Calretinin | MilliporeSigma, MAB1568 | ACs, RGCs | P10, P21 |
| Rbpms | MilliporeSigma, AB1362 | RGCs | P21 |
| Glutamine Synthetase | BD Biosciences, 610517 | Müller glia | P21 |
| GFAP | MilliporeSigma, G3893 | Reactive gliosis | P21 |
| IBA1 | WakoFujifilm, 019-19741 | Microglia | P21 |
| p27<sup>Kip1</sup> | BD Biosciences, 610241 | Müller glia | P21 |
| P2Y12R | AnaSpec, 55043A | Microglia | P21 |
| CD68 | Abcam, 125212 | Macrophages | P21 |
| Conjugated lectin | Vector Laboratories, DL-1177-1 | Mainly vasculature | P21 |
| VEGFR2 | Cell Signaling Technology, 55B11 | Vasculogenesis | P14 |

OS=outer segment, HC=horizontal cell, AC=amacrine cell, RGC=retinal ganglion cell, BC=bipolar cell

Procedures of IHC staining with paraffin-embedded sections are outlined as follows. Slides were treated with hot citrate buffer for antigen retrieval and subsequently blocked with a blocking buffer of 5% donkey serum, 1% bovine serum albumin, 0.1% Triton X-100 in 1X phosphate-buffered saline (PBS) (pH 7.4) for 2 hours. Slides were then incubated with primary antibodies (listed in Table 1) at 4°C overnight. Slides were washed with 1X PBS containing 0.01% TritonX-100 (PBST) for 30 min and then incubated with specific secondary antibodies for 1 hour. Slides were washed and mounted with hard set mounting medium with DAPI (Vectashield, Vector Laboratories).

Procedures of IHC staining with cryosections are outlined as follows. Dissected eyes were fixed in 4% paraformaldehyde for 1 hour, followed by the removal of lens. Tissues were subsequently immersed in increasing concentrations of phosphate-buffered sucrose solutions, and incubated overnight at 4°C in 20% phosphate-buffered sucrose solution. Tissues were incubated in a 1:1 solution mixture of OCT (optimal cutting temperature; Sakura Finetek) and 20% phosphate-buffered sucrose for 1 hour and then embedded in OCT and snap frozen. Cryosections were cut by 10 microns per section on a Cryotome E (Thermo Fisher Scientific). Blocking and antibody staining were performed as described above.

For vascular immunostaining and imaging of vasculature, mice were anesthetized with a cocktail of ketamine (86.9 mg/kg) and xylazine (10 mg/kg), then cardiac perfused with ice cold PBS with heparin (3U/mL) prior to surgical dissection of eyes. Whole-amount staining of conjugated lectin was performed with P21 retinae. Retinal samples were fixed overnight in 4% paraformaldehyde, then blocked and stained.

All cross-section slides were imaged taken on a Leica DB5500 microscope. All images were acquired at 1000 μm from ONH for ≥ P10 samples and at 500 μm from ONH for < P10 samples.

For 3D reconstruction of retinal vasculature, lectin-positive vasculature and *pNrl-eGFP*-positive photoreceptor nuclei of the retina were visualized using confocal laser scanning microscopy (Zeiss LSM 800, 20x) and Z-stacks that encompassed full depth of the vascular network (IVP-DVP) and ONL (0.5µm slices). Vascular and photoreceptor elements captured within images were reconstructed using Imaris 9.9 software (Oxford Instruments). Following automatic surface renderings of vascular and photoreceptor elements, manual editing was performed to delete erroneous process segments.

### Cell Count Analysis

Number of fluorescent objects were tallied in defined ranges as described in figure legends. 5 biological replicates of each genotype were used in the statistical analysis. Pairwise t-test was performed with p < 0.05, CI:95% using Graphpad Prism 10 (GraphPad Software).

### Electroretinography (ERG)

Mice were dark-adapted overnight prior to ERG tests. ERG tests on 3MO mice were conducted on the same UTAS-E3000 Visual Electrodiagnostic System (LKC Technologies Inc) that was employed in the published study with 1MO mice^4^, according to the same setting and protocol used in the previous study^44^. In short, for dark-adapted responses, electrode measurements were recorded with full-field light flashes (10 μs) of increasing intensity, the maximum flash intensity was 0.895 cd*s/m^2^. For light-adapted responses, mice were exposed to 10 μs light flashes of increasing intensity, maximum flash intensity for light-adapted testing was 2.672 cd*s/m^2^. ERG responses of biological replicates were recorded, averaged, and analyzed using GraphPad Prism 10 (GraphPad Software). The mean peak amplitudes were plotted against log values of light intensities (cd*s/m^2^). The statistics were obtained by using analysis of variance (ANOVA) or pairwise t-test.

### Quantitative PCR (qPCR)

For P0, P3 and P5 samples, each RNA sample was extracted from 4 retinae of 2 mice using the NucleoSpin RNA Plus kit (Macherey-Nagel). For samples older than P10, each RNA sample was extracted from 2 retinae of a mouse. 2 μg of RNA was used to produce cDNA using First Strand cDNA Synthesis kit (Roche). Primers used in this study were listed in Table 2. The reaction master mix consisted of EvaGreen polymerase (Bio-Rad Laboratories), 1 μM primer mix, and diluted cDNA samples. Samples were run using a two-step 40-cycle protocol on a Bio-Rad CFX96 Thermal Cycler (Bio-Rad Laboratories). Pairwise t-test was performed with p < 0.05, CI:95% using Graphpad Prism 10 (GraphPad Software).

**Table 2.**
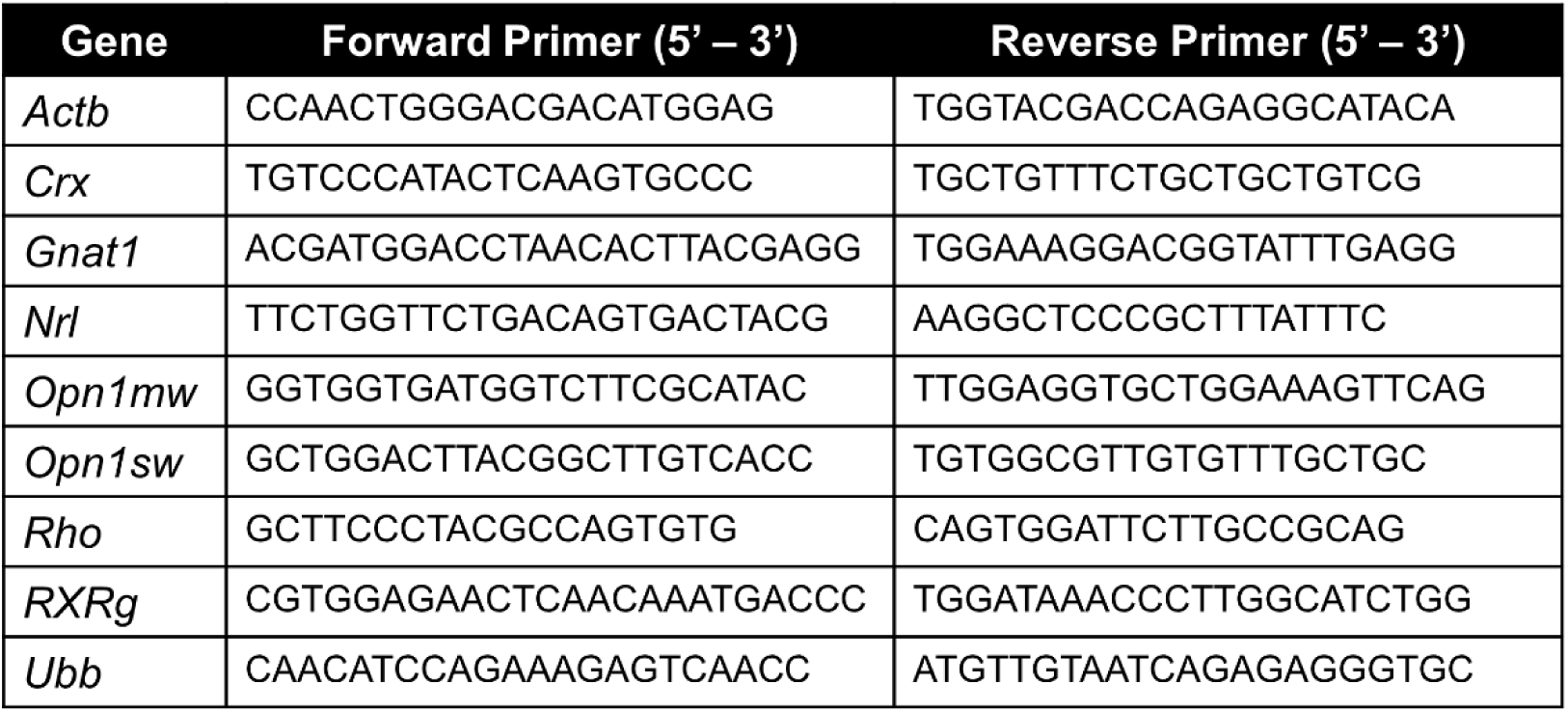
qPCR primers.

## ETHICS STATEMENT

The animal study was reviewed and approved by Washington University in St. Louis Institutional Animal Care and Use Committee.

## AUTHOR CONTRIBUTIONS

The principal investigators, SC and RSA, acquired the funding to support the project and oversaw the progress. CS performed IHC staining, imaging, ERG, qPCR and associated analysis. CP performed cardiac perfusion and whole-amount staining of conjugated lectin, 3D reconstruction imaging analysis. YZ assisted in sample collection. CS, CP and SC prepared the manuscript. All authors edited and revised the manuscript.

## ACKNOWLEDGMENTS

Authors are grateful to Ms. Mingyan Yang and Dr. Guangyi Ling for technical assistance. The following grants were involved in this study: Knights Templar Pediatric Ophthalmology Career-Starter Research Grant (CS), R01 EY012543 (SC), R01 EY032136 (SC), Widłak Family CRX Research Fund (SC), Jeffrey T. Fort Innovation Fund (RSA), Siteman Retina Research Grant (RSA), NIH P30 EY002687 (WUSTL DOVS), an unrestricted grant from Research to Prevent Blindness (WUSTL DOVS).

## REFERENCES

1 Sun, C. & Chen, S. Disease-causing mutations in genes encoding transcription factors critical for photoreceptor development. Frontiers in Molecular Neuroscience 16 (2023).

2 Tran, N. M. & Chen, S. Mechanisms of blindness: animal models provide insight into distinct CRX-associated retinopathies. Developmental dynamics : an official publication of the American Association of Anatomists 243, 1153–1166 (2014).

3 Furukawa, T., Morrow, E. M., Li, T., Davis, F. C. & Cepko, C. L. Retinopathy and attenuated circadian entrainment in Crx-deficient mice. Nature Genetics 23, 466 (1999).

4 Zheng, Y., Sun, C., Zhang, X., Ruzycki, P. A. & Chen, S. Missense mutations in CRX homeodomain cause dominant retinopathies through two distinct mechanisms. Elife 12, RP87147 (2023). PMC10645426.

5 Tran, N. M. et al. Mechanistically distinct mouse models for CRX-associated retinopathy. PLoS genetics 10, e1004111–e1004111 (2014).

6 Zheng, Y., Stormo, G. D. & Chen, S. Aberrant homeodomain-DNA cooperative dimerization underlies distinct developmental defects in two dominant CRX retinopathy models. Genome Res 35, 242–256 (2025). PMC11874979.

7 Freund, C. L. et al. Cone-Rod Dystrophy Due to Mutations in a Novel Photoreceptor-Specific Homeobox Gene (CRX) Essential for Maintenance of the Photoreceptor. Cell 91, 543–553 (1997).

8 Nichols II, L. L. et al. Two novel CRX mutant proteins causing autosomal dominant Leber congenital amaurosis interact differently with NRL. Human Mutation 31, E1472–E1483 (2010).

9 Swaroop, A. et al. Leber Congenital Amaurosis Caused by a Homozygous Mutation (R90W) in the Homeodomain of the Retinal Transcription Factor CRX: Direct Evidence for the Involvement of CRX in the Development of Photoreceptor Function. Human Molecular Genetics 8, 299–305 (1999).

10 Martínez-Gil, N. et al. Cellular and molecular alterations in neurons and glial cells in inherited retinal degeneration. Front Neuroanat 16, 984052 (2022). PMC9548552.

11 Ameri, H., Hong, A. T. & Chwa, J. Loss of Peripheral Retinal Vessels in Retinitis Pigmentosa. Ophthalmol Sci 5, 100767 (2025). PMC12032902.

12 Nguyen, M., Sullivan, J. & Shen, W. Retinal vascular remodeling in photoreceptor degenerative disease. Exp Eye Res 234, 109566 (2023).

13 Kumari, A. & Borooah, S. The Role of Microglia in Inherited Retinal Diseases. Adv Exp Med Biol 1415, 197–205 (2023).

14 Hippert, C. et al. Muller glia activation in response to inherited retinal degeneration is highly varied and disease-specific. PLoS One 10, e0120415 (2015). PMC4368159.

15 Young, R. W. Cell proliferation during postnatal development of the retina in the mouse. Developmental Brain Research 21, 229–239 (1985).

16 Cao, Y. et al. Mechanism for Selective Synaptic Wiring of Rod Photoreceptors into the Retinal Circuitry and Its Role in Vision. Neuron 87, 1248–1260 (2015). PMC4583715.

17 Fu, Y. & Yau, K. W. Phototransduction in mouse rods and cones. Pflugers Arch 454, 805–819 (2007). PMC2877390.

18 Nikonov, S. S. et al. Mouse Cones Require an Arrestin for Normal Inactivation of Phototransduction. Neuron 59, 462–474 (2008).

19 Roberts, M. R., Hendrickson, A., McGuire, C. R. & Reh, T. A. Retinoid X Receptor γ Is Necessary to Establish the S-opsin Gradient in Cone Photoreceptors of the Developing Mouse Retina. Investigative Ophthalmology & Visual Science 46, 2897–2904 (2005).

20 Hoover, F., Seleiro, E. A., Kielland, A., Brickell, P. M. & Glover, J. C. Retinoid X receptor gamma gene transcripts are expressed by a subset of early generated retinal cells and eventually restricted to photoreceptors. J Comp Neurol 391, 204–213 (1998).

21 Fong, S. L. Characterization of the human rod transducin alpha-subunit gene. Nucleic Acids Res 20, 2865–2870 (1992). PMC336934.

22 Arshavsky, V. Y., Lamb, T. D. & Pugh, E. N. G Proteins and Phototransduction. Annual Review of Physiology 64, 153–187 (2002).

23 Sherry, D. M., Wang, M. M., Bates, J. & Frishman, L. J. Expression of vesicular glutamate transporter 1 in the mouse retina reveals temporal ordering in development of rod vs. cone and ON vs. OFF circuits. J Comp Neurol 465, 480–498 (2003).

24 Johnson, J. et al. Vesicular glutamate transporter 1 is required for photoreceptor synaptic signaling but not for intrinsic visual functions. J Neurosci 27, 7245–7255 (2007). PMC2443709.

25 Pasteels, B., Rogers, J., Blachier, F. & Pochet, R. Calbindin and calretinin localization in retina from different species. Vis Neurosci 5, 1–16 (1990).

26 Greferath, U., Grünert, U. & Wässle, H. Rod bipolar cells in the mammalian retina show protein kinase C-like immunoreactivity. J Comp Neurol 301, 433–442 (1990).

27 Balasubramanian, R. & Gan, L. Development of Retinal Amacrine Cells and Their Dendritic Stratification. Curr Ophthalmol Rep 2, 100–106 (2014). PMC4142557.

28 Rodriguez, A. R., de Sevilla Müller, L. P. & Brecha, N. C. The RNA binding protein RBPMS is a selective marker of ganglion cells in the mammalian retina. J Comp Neurol 522, 1411–1443 (2014). PMC3959221.

29 Poitry, S., Poitry-Yamate, C., Ueberfeld, J., MacLeish, P. R. & Tsacopoulos, M. Mechanisms of glutamate metabolic signaling in retinal glial (Müller) cells. J Neurosci 20, 1809–1821 (2000). PMC6772920.

30 Levine, E. M., Close, J., Fero, M., Ostrovsky, A. & Reh, T. A. p27Kip1 Regulates Cell Cycle Withdrawal of Late Multipotent Progenitor Cells in the Mammalian Retina. Developmental Biology 219, 299–314 (2000).

31 Guo, L., Choi, S., Bikkannavar, P. & Cordeiro, M. F. Microglia: Key Players in Retinal Ageing and Neurodegeneration. Front Cell Neurosci 16, 804782 (2022). PMC8968040.

32 Fan, W. et al. Retinal microglia: Functions and diseases. Immunology 166, 268–286 (2022).

33 Fruttiger, M. Development of the Mouse Retinal Vasculature: Angiogenesis Versus Vasculogenesis. Investigative Ophthalmology & Visual Science 43, 522–527 (2002).

34 Stahl, A. et al. The Mouse Retina as an Angiogenesis Model. Investigative Ophthalmology & Visual Science 51, 2813–2826 (2010).

35 Wang, X., Bove, A. M., Simone, G. & Ma, B. Molecular Bases of VEGFR-2-Mediated Physiological Function and Pathological Role. Frontiers in Cell and Developmental Biology Volume 8 - 2020 (2020).

36 Okabe, K. et al. Neurons Limit Angiogenesis by Titrating VEGF in Retina. Cell 159, 584–596 (2014).

37 Sun, C. & Chen, S. Gene Augmentation for Autosomal Dominant CRX-Associated Retinopathies. Adv Exp Med Biol 1415, 135–141 (2023). PMC11010719.

38 Nikonov, S. S. et al. Photoreceptors of Nrl -/- mice coexpress functional S- and M-cone opsins having distinct inactivation mechanisms. J Gen Physiol 125, 287–304 (2005). PMC2234018.

39 Haider, N. B., Naggert, J. K. & Nishina, P. M. Excess cone cell proliferation due to lack of a functional NR2E3 causes retinal dysplasia and degeneration in rd7/rd7 mice. Hum Mol Genet 10, 1619–1626 (2001).

40 Hu, W. et al. Expression of VLDLR in the Retina and Evolution of Subretinal Neovascularization in the Knockout Mouse Model’s Retinal Angiomatous Proliferation. Investigative Ophthalmology & Visual Science 49, 407–415 (2008).

41 Daich Varela, M., et al. Coats-like Vasculopathy in Inherited Retinal Disease: Prevalence, Characteristics, Genetics, and Management. Ophthalmology 130, 1327–1335 (2023). PMC10937259.

42 Heath Jeffery, R. C. & Chen, F. K. Macular neovascularization in inherited retinal diseases: A review. Survey of Ophthalmology 69, 1–23 (2024).

43 Akimoto, M. et al. Targeting of GFP to newborn rods by Nrl promoter and temporal expression profiling of flow-sorted photoreceptors. Proc Natl Acad Sci U S A 103, 3890–3895 (2006). PMC1383502.

44 Sun, C., Zhang, X., Ruzycki, P. A. & Chen, S. Essential Functions of MLL1 and MLL2 in Retinal Development and Cone Cell Maintenance. Front Cell Dev Biol 10, 829536 (2022). PMC8864151.

