## Supplemental Figure 1-12 for "Aberrant retinal structure and vasculature in mouse models of dominant retinopathies caused by *CRX* homeodomain mutations": Aberrant retinal structure and vasculature in mouse models of dominant retinopathies caused by CRX homeodomain mutations Supplemental Figures.pdf

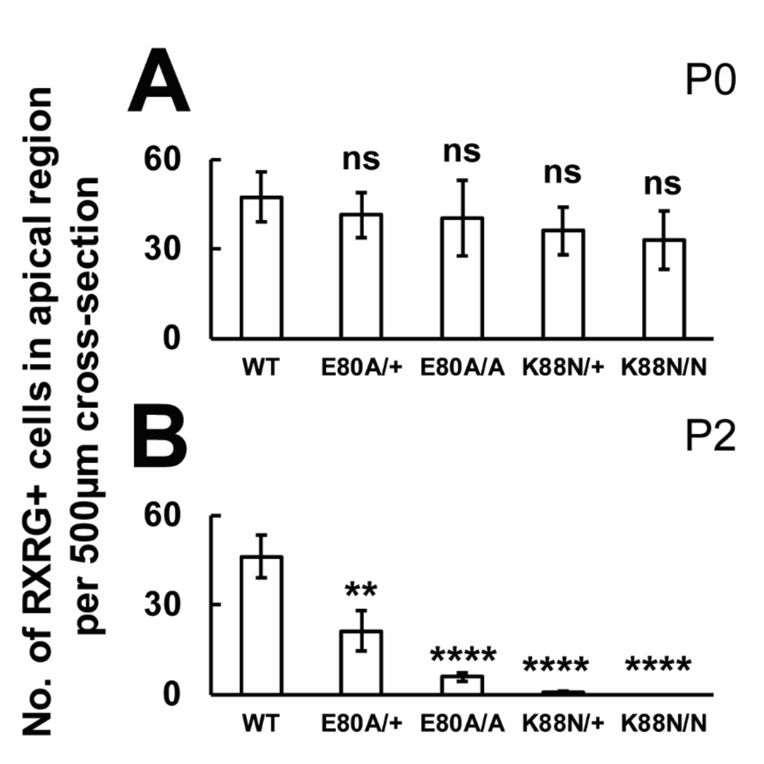


Supplemental Figure 1.

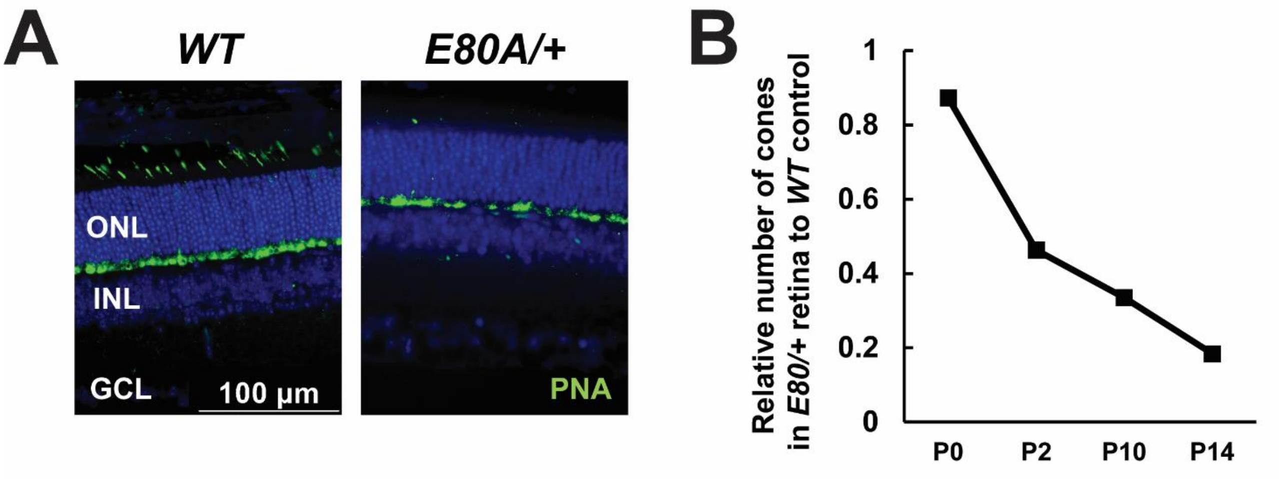


Supplemental Figure 2.

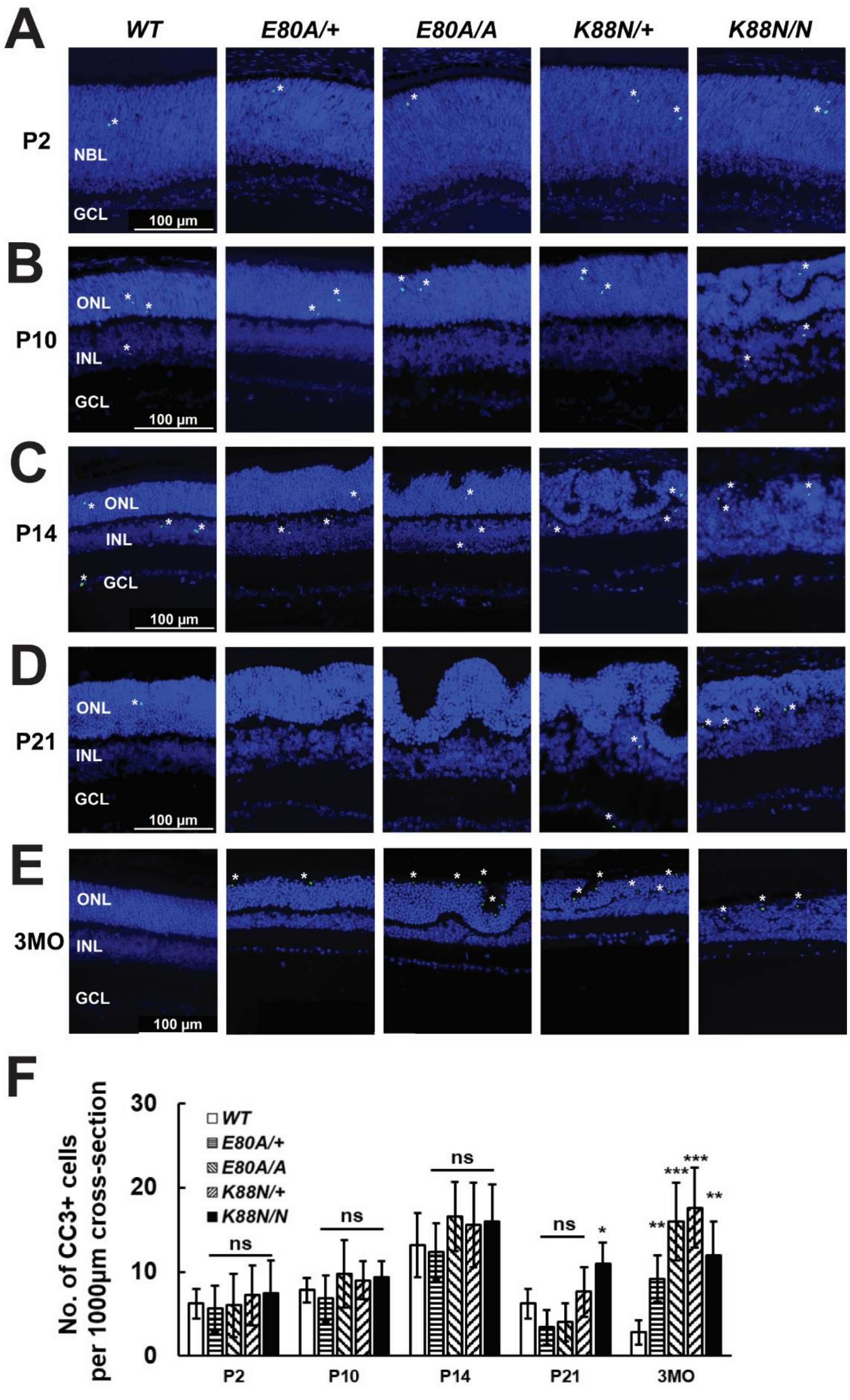


Supplemental Figure 3.

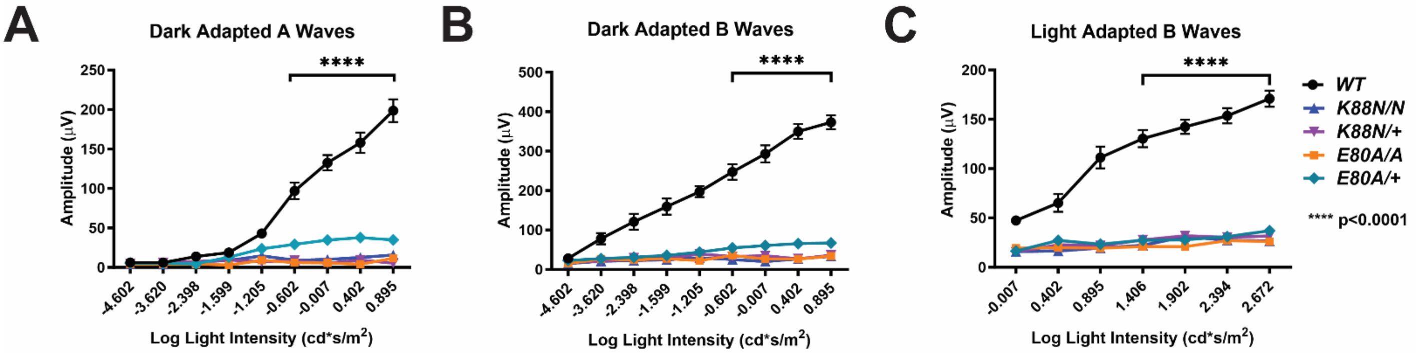


**D**

| Reduction % in E80A/+ ERG at age | Dark-adapted A-waves | | Dark-adapted B-waves | |
|---|---|---|---|---|
| | 0.402 cd*s/m$^2$ Mean (SD) | 0.895 cd*s/m$^2$ Mean (SD) | 0.402 cd*s/m$^2$ Mean (SD) | 0.895 cd*s/m$^2$ Mean (SD) |
| 1MO | 51.4 (16.1) | 40.9 (10.2) | 63.8 (14.8) | 61.7 (10.8) |
| 3MO | 76.0 (2.4) | 82.5 (2.0) | 81.2 (1.5) | 79.4 (0.7) |

Supplemental Figure 4.

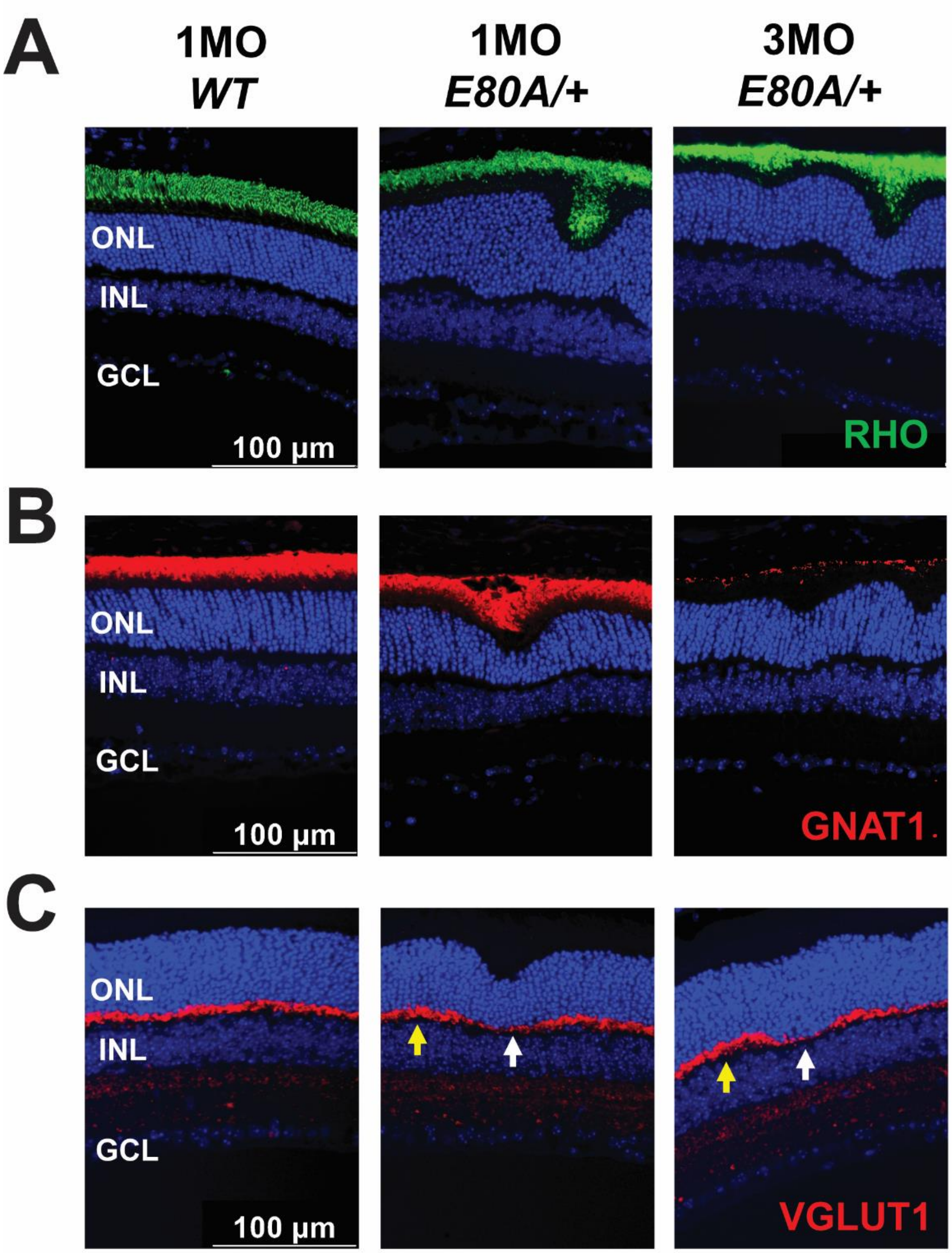


Supplemental Figure 5.

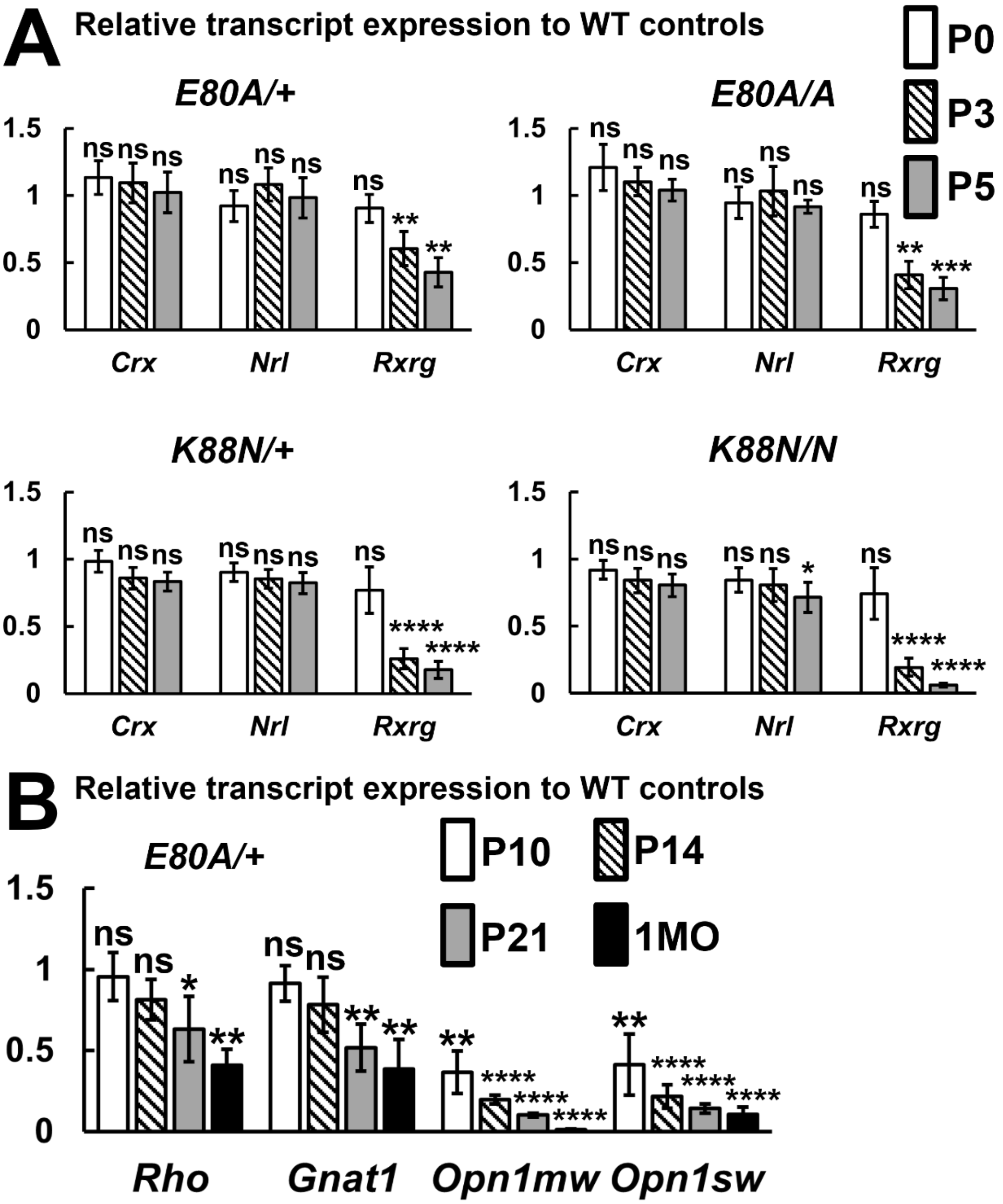


Supplemental Figure 6.

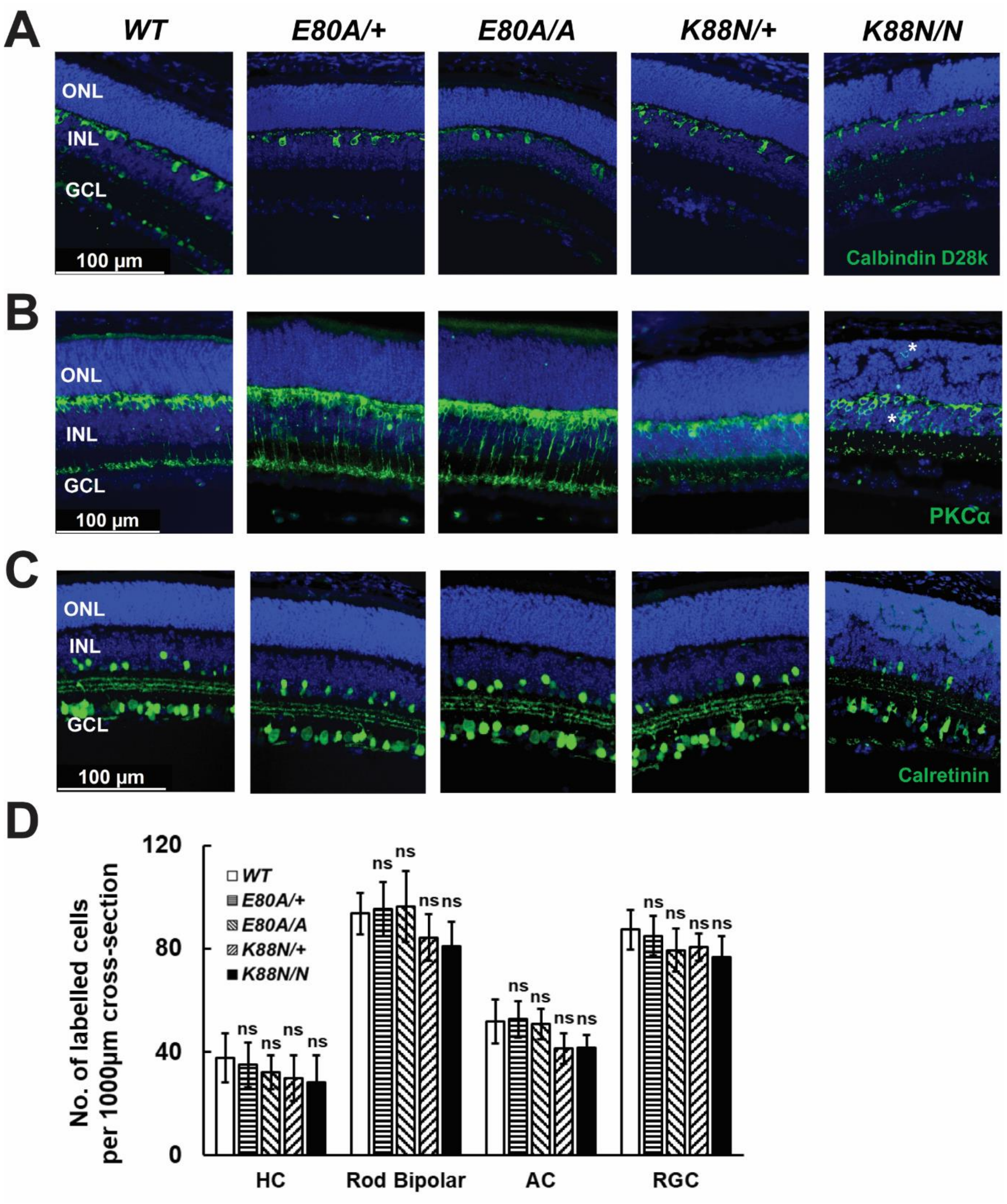


Supplemental Figure 7.

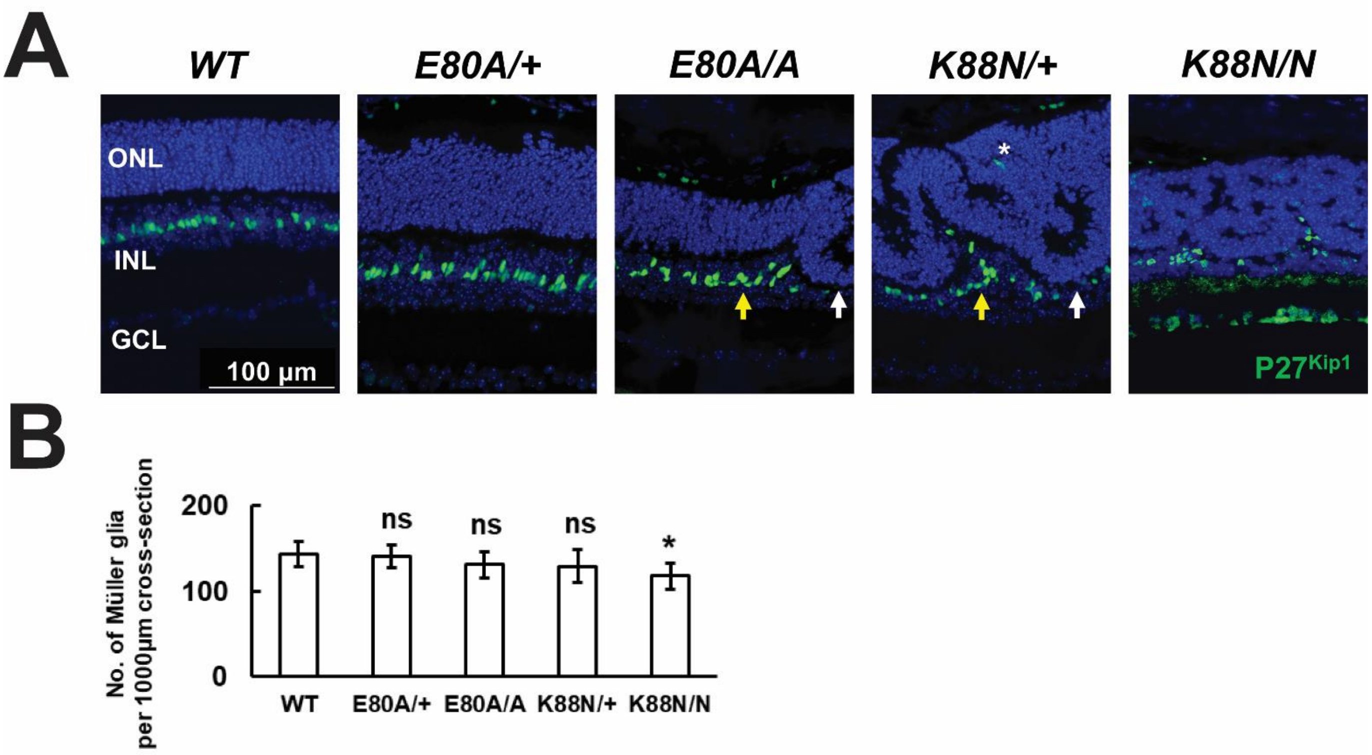


Supplemental Figure 8.

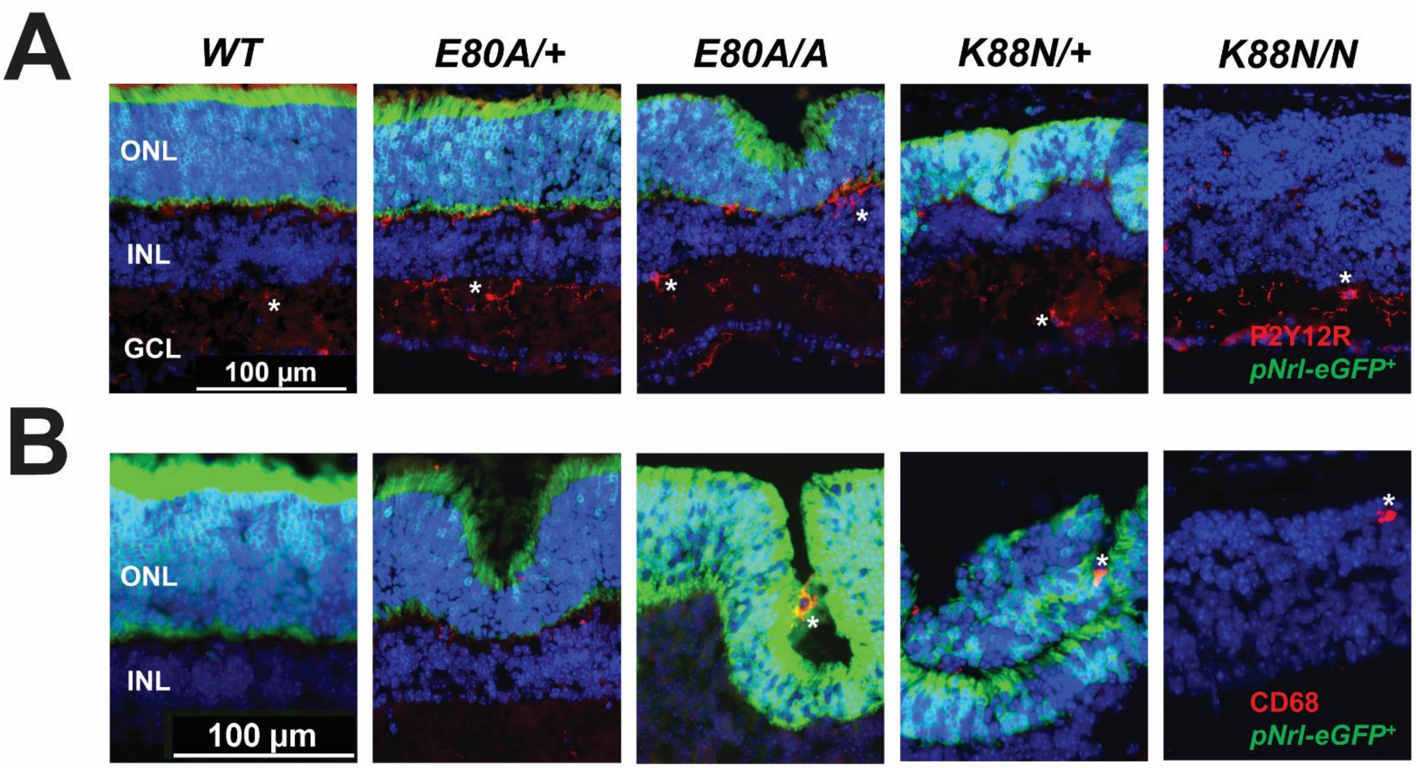


Supplemental Figure 9.

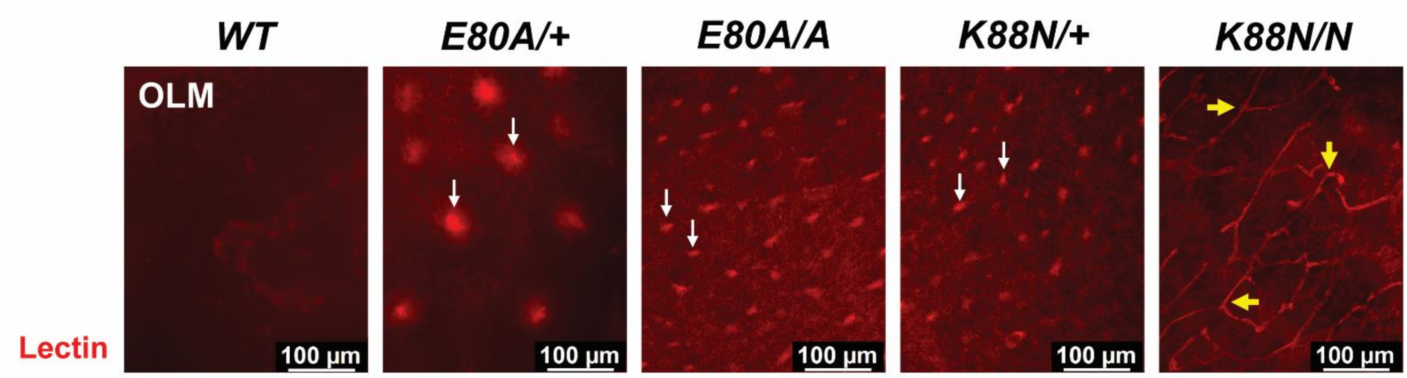


Supplemental Figure 10.

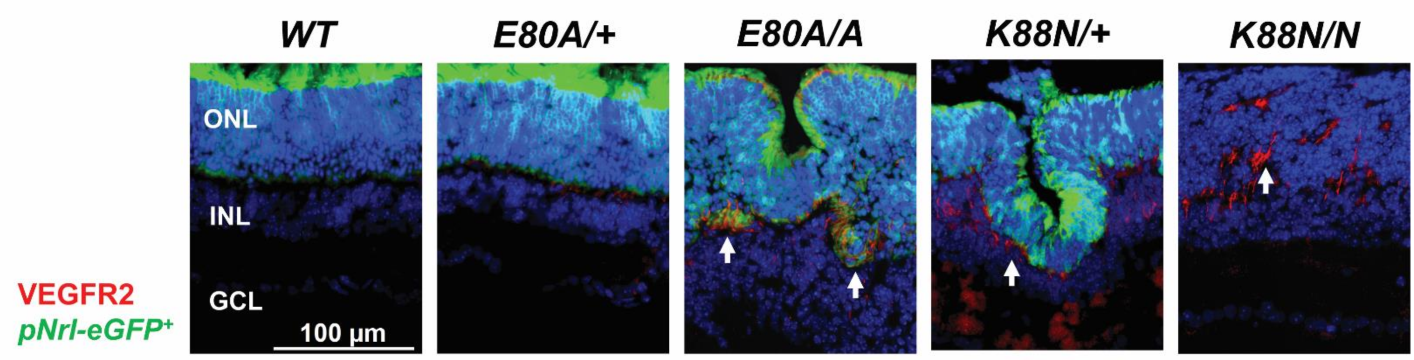


Supplemental Figure 11.

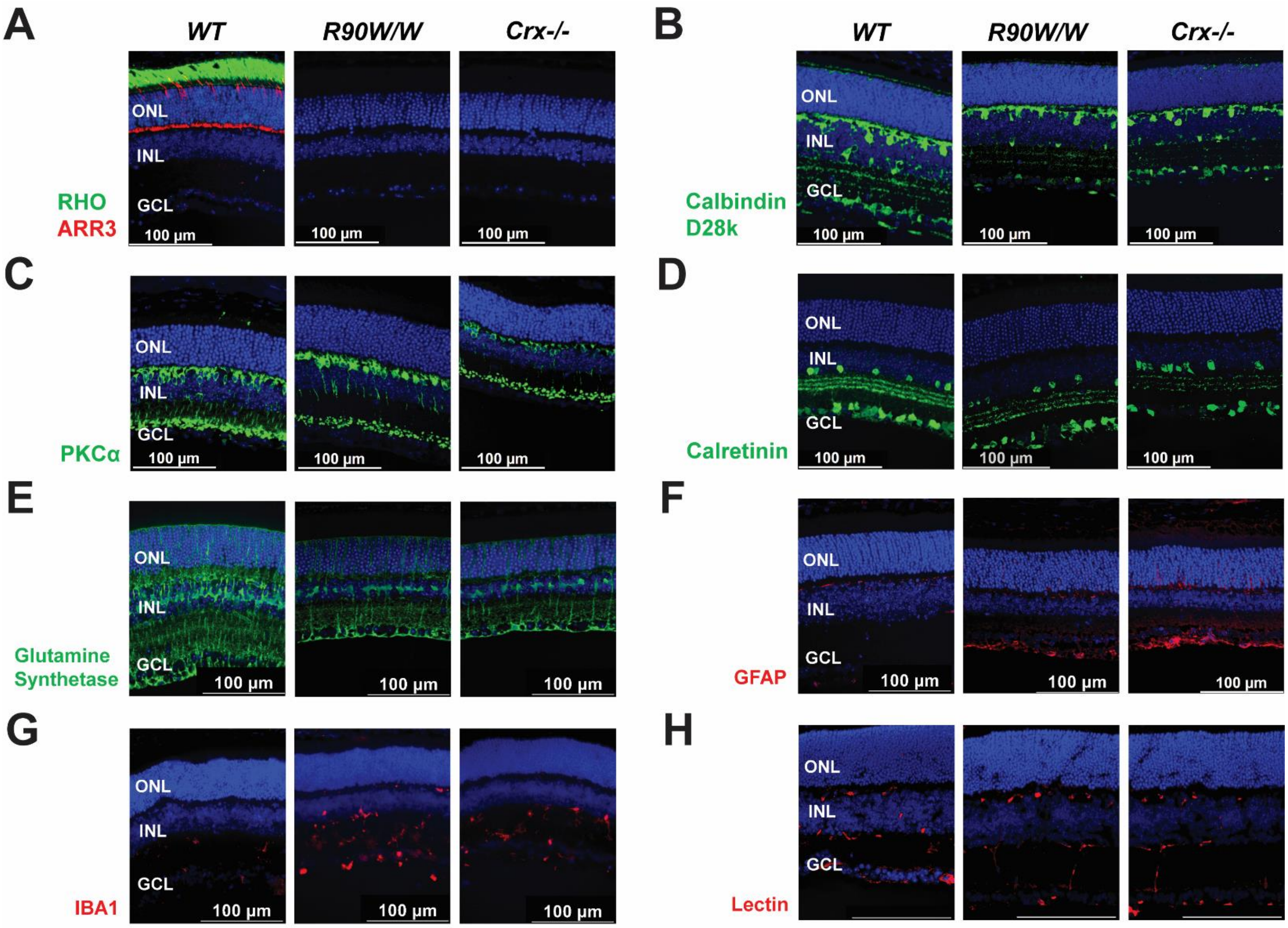


Supplemental Figure 12.

**SUPPLEMENTAL FIGURE LEGENDS**

Supplemental Figure 1. (A) Cell counts for RXRG+ cells between 500µm and 1000µm cross-sections of (A) P0, (B) P2 *WT* and mutant retinae. Error bars represent SD on mean values (n=4). Statistical analysis by pairwise t-test happens between *WT* and each mutant group. Asterisks (**, *****) denote p≤0.05 and p≤0.0001 respectively, ns means not significant.

Supplemental Figure 2. (A) IHC staining of conjugated peanut agglutinin (PNA, in green) in P14 *WT* and $Crx^{E80A/+}$ retinae. Nuclei are stained by DAPI (in blue). Scale bar = 100µm. (B) Relative numbers of cones in $Crx^{E80A/+}$ retinae to *WT* controls. Cell counts (n=4) at P0 and P2 are based IHC staining of RXRG. Cell counts (n=4) at P10 and P14 are based on IHC staining of ARR3.

Supplemental Figure 3. IHC staining of cleaved caspase 3 (CC3, in green) in (A) P2, (B) P10, (C) P14, (D) P21, (E) 3MO *WT* and mutant retinae. Asterisks indicate examples of CC3+ cells. Nuclei are stained by DAPI (in blue). Scale bar = 100µm. (F) Cell counts for CC3+ cells between 500µm and 1500µm cross-sections of *WT* and mutant retinae. Error bars represent SD on mean values (n=6). Statistical analysis by pairwise t-test happens between *WT* and each mutant group. Asterisks (*, **, ***, ****) denote p≤0.05, p≤0.01, p≤0.001, and p≤0.0001, ns means not significant.

Supplemental Figure 4. (A) Dark adapted A waves, (B) Dark adapted B waves, (C) light adapted B waves of ERG responses in 3MO *WT* and mutant retinae. Error bars represent SD on mean values (n=6). Statistical analysis is performed by ANOVA. Asterisks (*****) denote p≤0.0001, representing statistical significance between *WT* and each mutant group. (D) Reduction in dark-adapted A and B waves of ERG responses (two highest light intensities) in $Crx^{E80A/+}$ retinae as compared to those of *WT* controls at 1 and 3MO. SD is calculated on mean values (n=6).

Supplemental Figure 5. IHC staining of (A) RHO (in green), (B) GNAT1 (in red), (C) VGLUT1 (in red) in 1MO *WT*, 1MO $Crx^{E80A/+}$, and 3MO $Crx^{E80A/+}$ retinae. Nuclei are stained by DAPI (in blue). Scale bar = 100µm. White arrows in (C) indicate regions with diminished labelling intensity. Yellow arrows in (C) indicate regions with abundant labelling intensity.

Supplemental Figure 6. qPCR analysis on selected genes in (A) neonatal $Crx^{E80A}$ and $Crx^{K88N}$ mutant retinae, (B) $Crx^{E80A/+}$ retinae at various ages. Relative expression is normalized to age-matched *WT* controls. Error bars represent SD on mean values (n=4). Statistical analysis by pairwise t-test happens between *WT* and each mutant group. Asterisks (*, **, ***, ****) denote p≤0.05, p≤0.01, p≤0.001, and p≤0.0001, ns means not significant.

Supplemental Figure 7. IHC staining of (A) calbindin D28K (in green), (B) PKCα (in green), (C) calretinin (in green) in P10 *WT* and mutant retinae. Asterisks indicate mislocalized cells in $Crx^{K88N/N}$ retinae. Nuclei are stained by DAPI (in blue). Scale bar = 100µm. (D) Cell counts for different retinal cell types between 500µm and 1500µm cross-sections of P10 *WT* and mutant retinae. Error bars represent SD on mean values (n=4). Statistical analysis by pairwise t-test happens between *WT* and each mutant group. ns means not significant.

Supplemental Figure 8. (A) IHC staining of $P27^{Kip1}$ (in green) in P21 *WT* and mutant retinae. White arrows indicate regions lacking labelled cells. Yellow arrows indicate clusters of labelled cells. Asterisk indicates an example of mislocalized cells. Nuclei are stained by DAPI (in blue). Scale bar = 100µm. (B) Cell counts for $P27^{Kip1}$ labelled Müller glia between 500µm and 1500µm cross-sections of P21 *WT* and mutant retinae. Error bars represent SD on mean values (n=4). Statistical analysis by pairwise t-test happens between *WT* and each mutant group. Asterisk (*) denotes p≤0.05, ns means not significant.

Supplemental Figure 9. IHC staining of (A) P2Y12R (in red), (B) CD68 (in red) in P21 *WT* and mutant retinae. Green cells are labelled by *pNrl-eGFP*. Asterisks indicate examples of labelled cells. Nuclei are stained by DAPI (in blue). Scale bar = 100µm.

Supplemental Figure 10. Whole-amount staining of conjugated lectin in P21 *WT* and mutant retinae. Images are taken at the outer limiting membrane (OLM). White arrows indicate examples of retinal rosettes. Yellow arrows indicate examples of vessels at OLM of *Crx*$^{K88N/N}$ retinae. Scale bar = 100μm.

Supplemental Figure 11. IHC staining of VEGFR2 (in red) in P14 *WT* and mutant retinae. Green cells are labelled by *pNrl-eGFP*. White arrows indicate regions with abundant labelling intensity. Nuclei are stained by DAPI (in blue). Scale bar = 100μm.

Supplemental Figure 12. IHC staining of (A) RHO (in green) and ARR3 (in red), (B) calbindin D28K (in green), (C) PKCα (in green), (D) calretinin (in green), (E) glutamine synthetase (in green), (F) GFAP (in red), (G) IBA1 (in red), (H) conjugated lectin (in red) in P21 *WT* and mutant retinae. Nuclei are stained by DAPI (in blue). Scale bar = 100μm.
